# Peptide abundance correlations in metaproteomics enhance taxonomic and functional analysis of the human gut microbiome

**DOI:** 10.1101/2025.03.13.642886

**Authors:** Zhongzhi Sun, Zhibin Ning, Qing Wu, Leyuan Li, Andrew C. Doxey, Daniel Figeys

## Abstract

Mass spectrometry (MS)-based proteomics is widely used for quantitative protein profiling and has become a powerful tool for studying protein interactions. However, most current research focuses on single-species proteomics to study protein interactions. Protein interactions within more complex microbiomes, composed of 100’s of bacterial species, remain largely unexplored. The human gut microbiome, closely linked to human health and disease, has become a key area of study using metaproteomics. Yet, due to the complexity of the microbiome, the interactions between gut microbes and their host remain largely unknown. In this study, we analyzed peptide abundance correlations within a metaproteomics dataset derived from *in vitro* cultured human gut microbiomes subjected to various drug treatments. Our analysis revealed that peptides from the same protein or taxon exhibited correlated abundance changes. By using t-SNE for visualization, we generated a peptide correlation map in which peptides from the same taxon formed distinct clusters. Furthermore, peptide abundance correlations enabled genome-level taxonomic assignments for a greater number of peptides. In single-species subsets of the dataset, peptide correlation networks constructed using taxon-based normalized peptide abundance (TNPA) linked peptides from functionally related proteins. These networks also provided insights into the potential functions of previously uncharacterized proteins. Altogether, our study demonstrates that analyzing peptide abundance correlations enhances both taxonomic and functional analyses in human gut metaproteomics research.

## Introduction

Despite many studies linking the human gut microbiome with human health and disease^1–5^, elucidating the mechanisms underlying the impact of the microbiome on host biology remains challenging, largely due to the complexity and the dark matter in gut microbial communities^6,7^. Different omics technologies, including metagenomics, metatranscriptomics, metaproteomics, and metabolomics, have become important tools for studying the human gut microbiome ^8^. One of these techniques, metaproteomics, measures the presence and abundance of proteins that drive biological functions within microbial communities, providing a direct glimpse of gut microbiome pathways ^9^.

Proteins involved in the same biological pathways tend to be co-regulated^10^. These functionally related proteins typically have similar expression profiles, leading to positive abundance correlations. Conversely, proteins involved in mutual inhibition, feedback regulation, or competition for binding sites tend to display negative abundance correlations^11,12^. Given accurate quantification using mass spectrometry, abundance correlations between proteins could reveal functional linkages, even for proteins that do not physically interact or colocalize, thus enabling the prediction of unknown protein functions ^13^. Protein abundance correlations have been applied to predict the functions of uncharacterized proteins in various model organisms, such as *Escherichia coli* ^14^, *Saccharomyces cerevisiae* ^15^, and *Homo sapiens* ^13^. However, these studies have been largely limited to single-species proteomics. In complex microbial communities, like the gut microbiome, protein abundance correlations remain unexplored. A previous study applied gene abundance correlation in metagenomics to predict microbial functional organization ^16^, but since proteins directly execute biological processes, studying protein abundance correlations is expected to more faithfully reflect functional dynamics in microbial communities. Given that over 40% of proteins from the human gut microbiome remain functionally uncharacterized ^17^, and that some of these unknown proteins have been implicated in disease development ^18,19^, understanding their functions is critical for advancing our knowledge of host-microbiome interactions.

Studying protein abundance correlations is more challenging in metaproteomics compared to single-species proteomics due to the difficulties in protein inference ^20^. In complex microbial communities, peptides can be shared among homologous proteins from different species, making it hard to assign them to their actual proteins of origin. Our previous studies showed that a large number of identified peptides were shared by multiple species, complicating taxonomic and protein source assignments ^21^. Moreover, even after protein inference, it remains debatable whether it is reasonable to aggregate peptide quantities into a single protein quantity, as peptides from the same protein-coding sequence could express different quantitative responses ^22^. Recently, peptide-centric metaproteomics analysis, which directly links peptides to their taxonomic and functional annotations, provides an alternative approach ^23^. Since mass spectrometry directly measures peptides instead of proteins, peptide-centric analysis is inherently reasonable and has been shown to be more sensitive and uncover features that are masked at the protein level ^24^.

Given the challenges in protein inference and the inherent advantages of peptide-centric analysis, we focus on peptide abundance correlations in this study. By leveraging peptide-centric analysis, we aim to overcome the biases associated with protein inference and provide a more accurate representation of functional and taxonomic relationships in microbial communities.

In this study, we analyzed peptide abundance correlations in a metaproteomics dataset derived from *in vitro* cultured human gut microbiomes subjected to different drug treatments, also referred to as perturbations. We explored the biological foundations of these correlations and visualized them using a peptide correlation map. Additionally, we applied peptide abundance correlations to improve the taxonomic assignment of peptides. Focusing on single-species subsets, we calculated taxon-based normalized peptide abundance (TNPA) and constructed peptide abundance correlation networks. These networks revealed functional linkages, providing new insights into the roles of previously uncharacterized microbial proteins.

## Methods

### Metaproteomics dataset

In this study, we analyzed a metaproteomics dataset comprising 672 raw files from human gut microbiomes of six individuals subjected to various drug treatments (112 samples per individual). Specifically, stool samples were treated with 109 different compounds—107 drugs, two DMSO samples as negative controls, and three kestose samples as positive controls (detailed in Supplementary Table S1)—and then *in vitro* cultured using the RapidAIM assay ^25^, a culture- and metaproteomics-based rapid method for studying individual microbiome responses to drugs. Detailed information on sample collection, preparation, and LC-MS/MS analysis has been documented in a separate study ^26^. Metaproteomics raw files obtained from LC-MS/MS were searched against the IGC (Integrated Gene Catalog) database ^27^ with MetaLab2.3 ^28^ with the MaxQuant ^29^ workflow, using default settings. Peptide quantification results were extracted from the “peptides.txt” file generated by MetaLab for subsequent analysis.

### Peptide abundance correlation calculation

**Pre-processing and peptide filtering.** Peptide abundance correlations across different perturbations were calculated separately for each individual. First, peptide identification and quantification results were extracted individually. Peptides were filtered by retaining only those with non-zero intensity in at least 20% of the samples for each individual (≥ 23 samples). The quantification results of peptides that passed this filtering step were used for further analysis.

**Peptide abundance log2-fold change (log2-FC) calculation.** Peptide raw intensity values were log2-transformed (with zeros replaced by 1 to enable proper transformation). The log2-fold change in peptide abundance for each sample was calculated by dividing the peptide log2-intensity under each treatment by the average log2-intensity of the two DMSO-treated control samples. For this calculation, zero values in the log2-transformed intensity were replaced with 1 to ensure proper log2-FC computation.

**Spearman correlation coefficients (SCC) calculation.** Peptide abundance correlations across different treatments were calculated for all peptide pairs using Spearman correlation coefficients (SCCs) of the log2-FC values across the treatments. The SCCs were computed using the *cor* function in R.

### Peptide annotations

To enable meaningful biological analysis, peptides were assigned to their protein sources and annotated with taxonomic and functional information using the following procedures.

**Protein source refinement.** We first generated a genome-level taxonomic profile for each individual to refine peptide protein source assignments and improve peptide annotation resolution. All identified peptides from each individual were mapped to their taxonomic sources by aligning them against MetaPep ^21^ records. Genome-distinct peptides, defined as peptides exclusively present in a single bacterial genome, were utilized to enhance the accuracy of assignments. In the subsequent step, only genomes with at least three genome-distinct peptides identified across all samples were considered for peptide protein source assignments.

**Peptide protein sources annotation.** Next, for peptide protein source assignments, protein sequences from the 4,744 UHGG representative genomes ^17^ were in-silico digested using DeepDigest ^30^ with the following parameters: miscleavage = 2, minimum peptide length = 7, maximum peptide length = 47 (default settings), and trypsin as the protease. Subsequently, all identified peptides were mapped to their unique protein source or multiple protein sources.

**Peptide taxonomic and functional sources annotation.** Each peptide was taxonomically annotated by assigning it to a specific genome source or the lowest common ancestor (LCA) of the genomes corresponding to all its protein sources. For functional annotations, protein functionally annotations were extracted from the UHGG ^17^ database, and each peptide was functionally annotated based on the functional annotation of its source protein(s).

### Visualize peptide abundance correlations with global peptide correlation maps

t-SNE (t-distributed stochastic neighbor embedding) was applied to visualize the peptide abundance correlations. The SCC matrix of peptide pairs from each individual was used as the input for the *Rtsne* package ^31^ with default parameters (perplexity = 30). In the peptide correlation map, each point represents a peptide, and the peptides were colored based on their taxonomic or functional annotations acquired as described in the previous section.

### Applying a machine learning model to predict peptide genome source

Genome-distinct peptides from the two genomes with the highest number of such peptides within the same bacteria family were extracted. Seventy percent of these peptides were randomly selected as the training dataset, while the remaining 30% constituted the test dataset. Additionally, genome-distinct peptides from other genomes within the same family and 500 randomly selected peptides from other families were included for further evaluation of the model’s performance.

For the training dataset, the Spearman correlation coefficients (SCCs) of each peptide’s abundance changes relative to all peptides in the dataset were calculated and used as input features. A Random Forest classifier was then trained using the *randomForest* package ^32^ in R with ntree = 500, importance = True. The model’s performance was assessed using confusion matrices calculated on two different combined test datasets: (1) genome-distinct peptides from the test dataset combined with genome-distinct peptides from other genomes of the same family, and (2) genome-distinct peptides from the test dataset combined with 500 randomly selected peptides from other families. Following these assessments, the trained model was used to predict genome sources of peptides from the same bacterial family that lacked genome-level taxonomic annotations.

### Single-species peptide abundance correlations

#### Calculation of taxon-based normalized peptide abundance (TNPA)

To mitigate the impact of changes in taxon abundance on peptide abundance and to better study the effect of different functions on peptide abundance, taxon-based normalized peptide abundance (TNPA) was calculated based on the original peptide abundance (OPA), total peptide abundance of all peptides from the same taxon in the sample (TPATS, Total Peptide Abundance per Taxon in a Sample), and an average level of peptide abundance (10^8^)using the following formula:

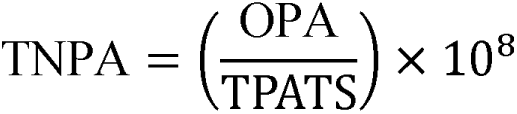

TNPA was calculated only for the top 10 genomes from each individual microbiome with the largest number of identified genome-distinct peptides. The subsequent single-species peptide abundance correlation analysis also focused on these genomes.

#### Single-species peptide abundance correlation network

Two types of single-species peptide abundance correlation networks were constructed using the SCCs of the log2-FC of OPA and the SCCs of log2-FC of TNPA. Peptide pairs with the top 5% SCCs were retained for network construction, which was carried out using the *igraph* package ^33^ in R. Nodes and edges from each network were extracted with the same package, and all peptides in networks were mapped to their protein source annotations.

Single-species peptide abundance correlation networks were visualized using *Gephi 0.10.1* with the Yifan Hu layout ^34^. Network modularity and clustering coefficients were calculated using the *modularity* and *transitivity* functions from the *igraph* package, respectively. Networks constructed with TNPA were used to study the correlations between peptides with related functions and to predict the functions of previously uncharacterized proteins.

### Data availability

Metaproteomics raw files used to compile analysis were deposited at the ProteomeXchange Consortium ^35^ via the PRIDE ^36^ partner repository as described in our previous study^26^. These files will be publicly available upon publication. Additionally, another dataset generated using a Q Exactive mass spectrometer is available with the data set identifiers PXD012724. All codes to perform the analysis in this study are available on GitHub at https://github.com/northomics/Peptide_Abundance_Correlations.

## Results

### Peptide abundance correlations in the metaproteomics dataset

Figure 1 summarizes the overall experimental design and research framework. Briefly, 6 individual microbiomes were individually exposed to 107 drugs and five controls using the RapidAIM assay ^25^ which is a 96-well plate assay that maintains the composition and function of the microbiome. The metaproteomes were extracted from each well and analyzed by metaproteomics. Peptide identification and quantification results were acquired as previously described ^26^. After acquiring peptide quantification results, peptides with non-zero values in ≥ 20% of samples from each individual were selected. Peptide abundance correlations for all peptide pairs from each individual were calculated using Spearman correlation coefficients (SCCs) for log2-transformed abundance fold changes from all 112 samples against the control group. Global peptide abundance correlation maps created with t-SNE of all six individuals colored with family-level taxonomic annotation of peptides showed a clear clustering of peptides from the same family (Figure 2A). In addition, these maps revealed substantial inter-individual variation in the taxonomic composition of microbiomes, which was further supported by peptide-based taxonomic profiling for each individual (Supplementary Figure 1). Among all six individuals, V52 has the largest number of quantified peptides (Figure 2B) and was selected as an example in the main text. For 21,363 peptides with non-zero values in ≥ 20% of samples from V52, SCC calculation yielded a total of 228,178,203 peptide pairs with an average SCC of 0.13 ± 0.35 (Figure 3A).

**Figure 1:**
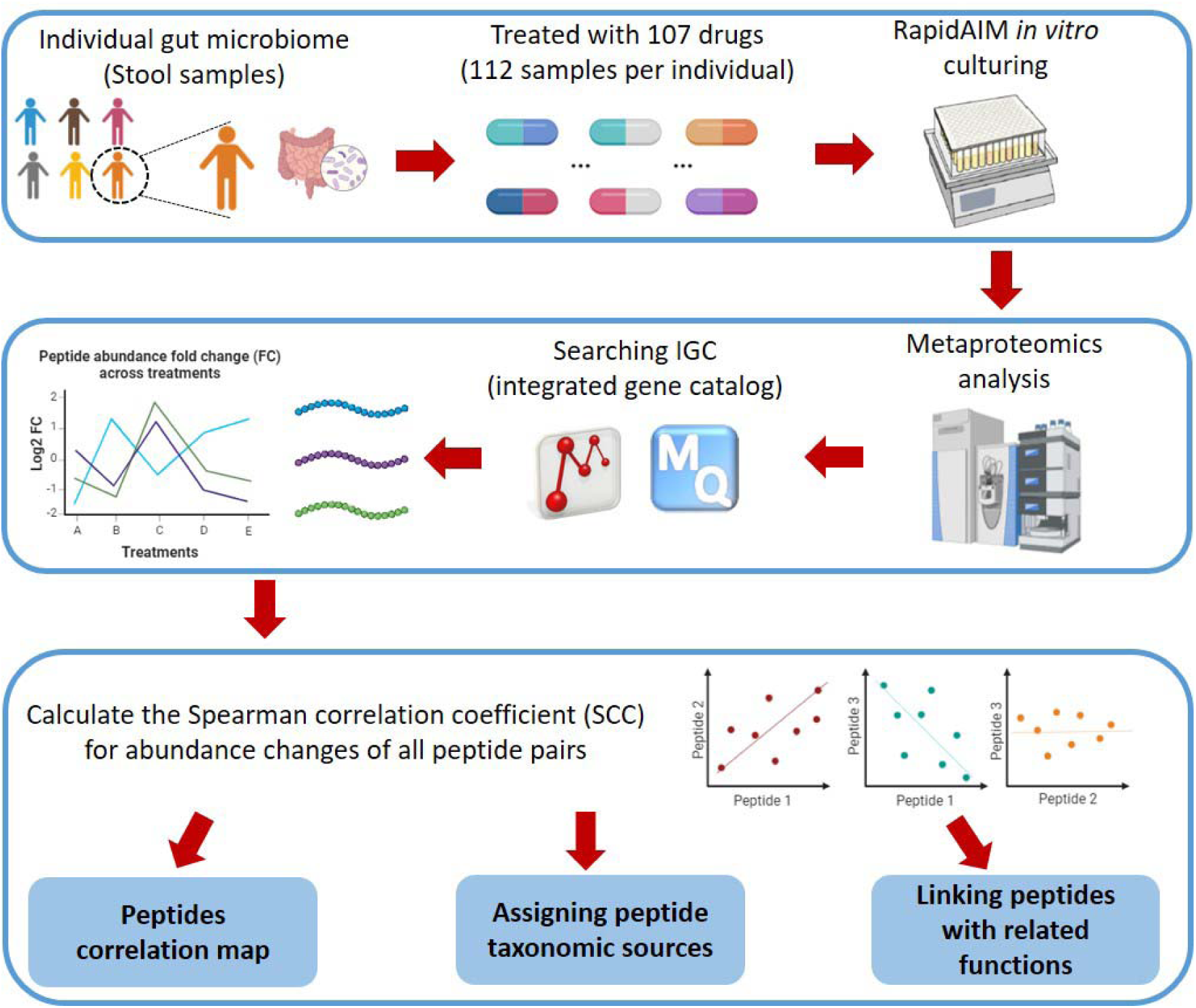
Experimental design and research framework. Gut microbiomes from six individuals were extracted from human stool samples. The microbiomes were then treated with 107 various drugs, resulting in 112 samples per individual, and *in vitro* cultured using RapidAIM. After culturing, samples were loaded to LC-MS/MS for metaproteomics analysis. Raw files were searched against the IGC database using the MaxQuant workflow from MetaLab V2.3, yielding a peptide abundance table. To measure peptide abundance correlations, we calculated the Spearman correlation coefficient (SCC) of peptide abundance fold changes relative to the control sample for all peptide pairs. We then created a peptide correlation map to visualize these correlations. Furthermore, we utilized these peptide abundance correlations to assign peptide taxonomic sources, connect peptides from functionally related proteins, and predict the functions of previously uncharacterized proteins.

**Figure 2:**
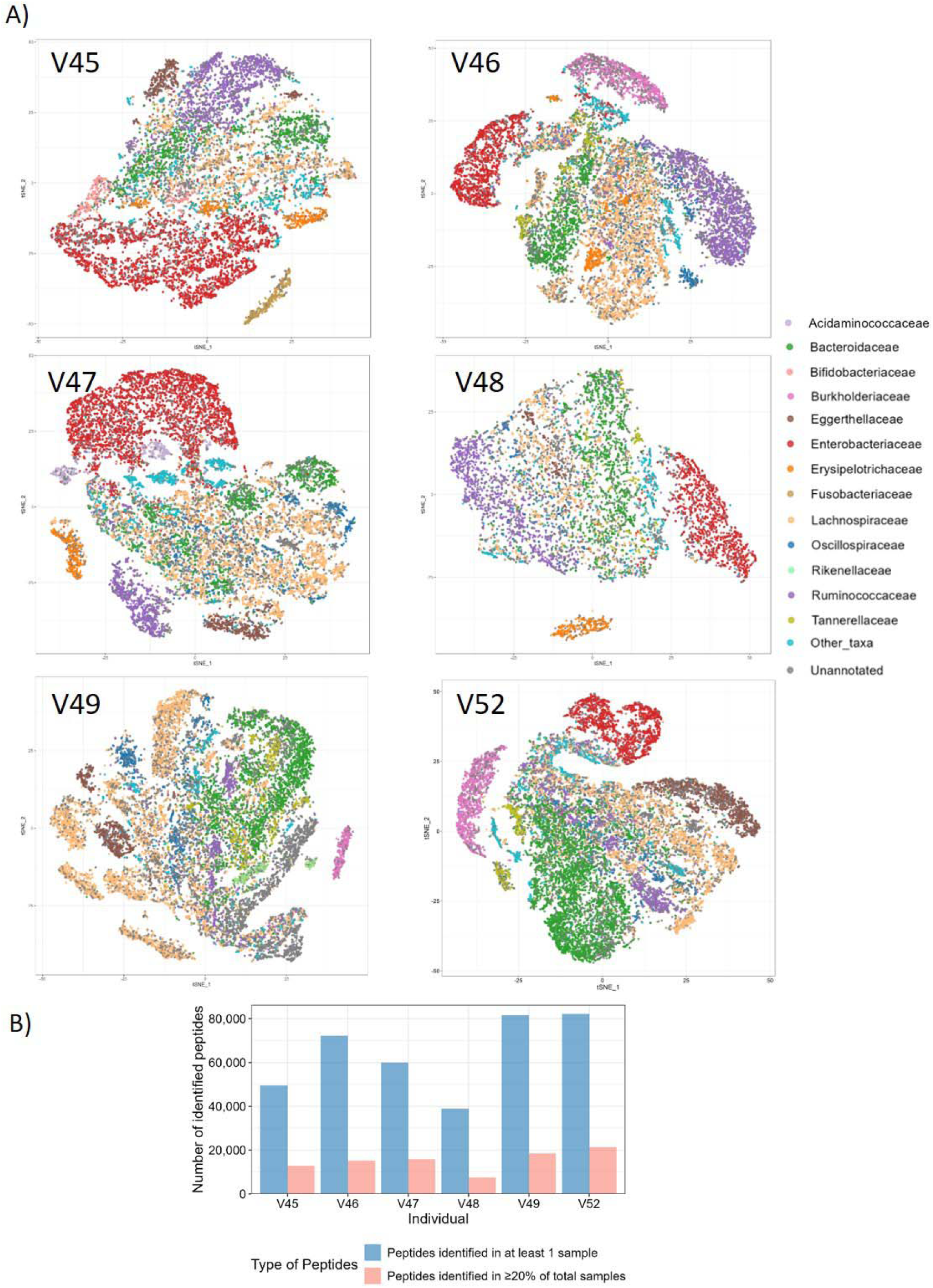
Peptide identification and abundance correlation across individuals. **A)** Peptide correlation maps generated using tSNE of all six individuals, colored by peptide family-level taxonomic annotations. **B)** Number of peptides identified from different individuals. For each individual, peptides identified in ≥ 20% of total samples were used for abundance correlation analysis and displayed in the peptide correlation map.

**Figure 3:**
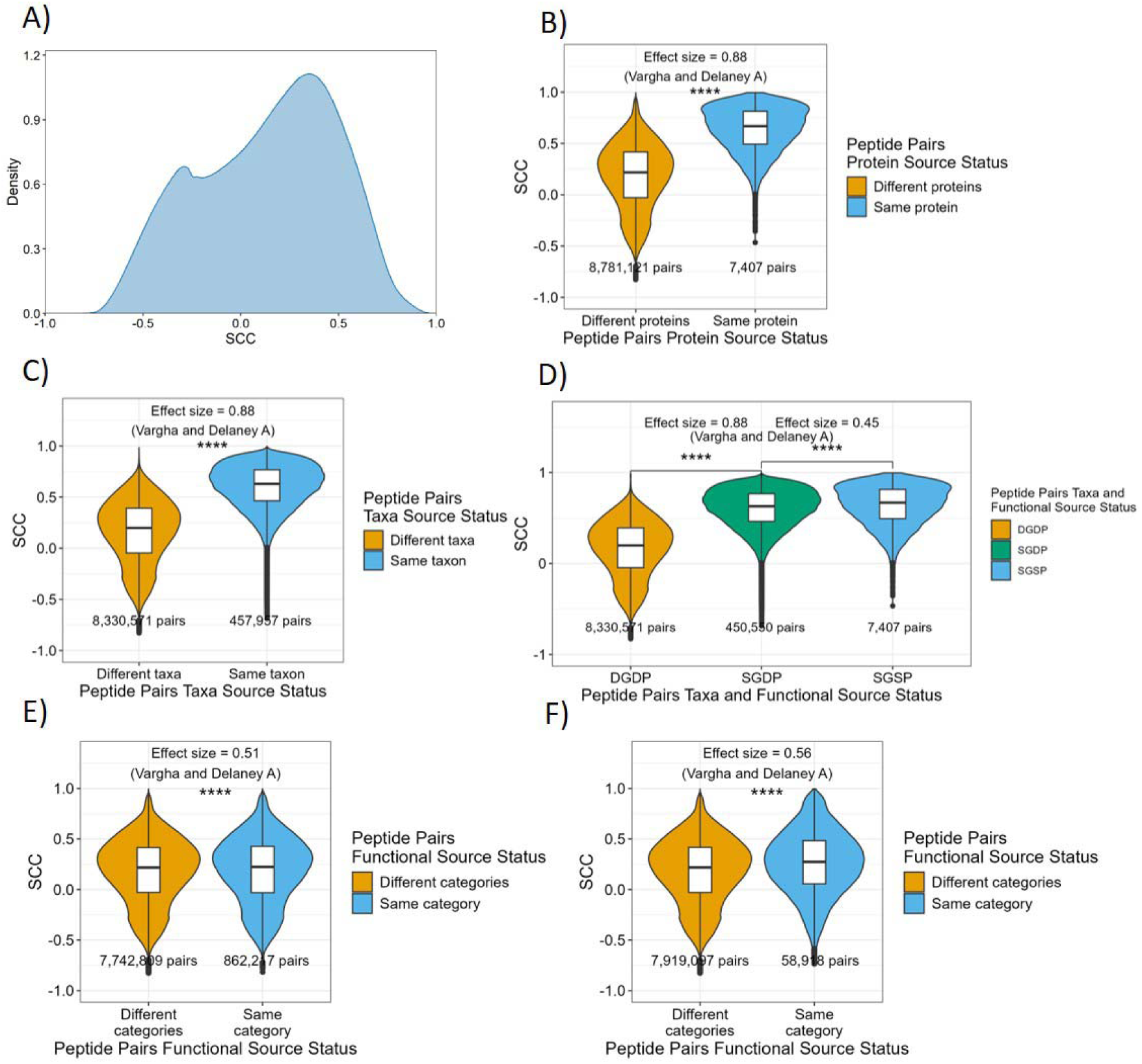
Peptide abundance correlations in the metaproteomics dataset of V52. **A)** Distribution of pairwise Spearman correlation coefficients (SCC) of peptide abundance fold changes for all peptide pairs. **B)** Comparison of the SCC of peptide pairs with both peptides from the same protein and peptide pairs of peptides from different proteins. **C)** Comparison of the SCC of peptide pairs of both peptides from the same taxon (genome) and peptide pairs of peptides from different taxa (genomes). **D)** Comparison of peptide pairs of both peptides from the same genome and same protein (SGSP), peptides from the same genome but different proteins (SGDP), and peptides from different genomes and different proteins (DGDP). **E)** Comparison of SCCs between peptide pairs from the same COG category and peptide pairs from different COG categories. **F)** Comparison of SCCs between peptide pairs from the same COG family and peptide pairs from different COG families. **** indicates statistical significance at the p ≤ 0.0001 levels by two-sided Mann-Whitney U test.

Upon assigning both protein sources and taxonomic sources to the peptides, we observed that peptides from the same protein and peptides from the same taxon exhibited higher abundance correlation. First, peptide pairs derived from the same protein (7,407 pairs) exhibited higher SCCs (0.63 ± 0.22) compared to peptide pairs from different proteins (8,781,121 pairs; SCC = 0.18 ± 0.32, with p ≤ 0.0001 by Mann-Whitney U test and a large effect size with Vargha and Delaney’s A of 0.88, Figure 3B). To be more intuitive, peptides from the same protein displayed more similar abundance profiles across samples than those from different proteins (Supplementary Figure 2). In addition, peptide pairs derived from the same genome (457,957) had an average SCC of 0.60±0.22, while those from different genomes (8,330,571) averaged an SCC of 0.16±0.31 (Figure 3C, p ≤ 0.0001 by Mann-Whitney U test and a large effect size with Vargha and Delaney’s A of 0.88). It is worth noting that even after excluding peptide pairs from the same protein, we found that peptides from different proteins within the same genome (450,550 pairs; SCC = 0.60 ± 0.22) still had higher SCCs than those from different genomes (8,330,571 pairs; SCC = 0.16 ± 0.31, p ≤ 0.0001 by Mann-Whitney U test, Vargha and Delaney’s A of 0.88), indicating that sourcing from the same genome contributes significantly to higher SCCs of peptide abundance changes (Figure 3D). We also assigned functional annotations to the peptides, however, the difference in SCCs between peptides from the same functional category and those from different functional categories was relatively minor at both high-level COG category (Figure 3E) and refined COG family level (Figure 3F), with negligible effect size, Vargha and Delaney’s A of 0.51 and 0.56, respectively.

Higher SCC of peptides from the same protein and peptides from the same taxon collectively suggested that studying peptide abundance correlations in metaproteomics datasets is biologically meaningful and can provide valuable insights into metaproteomics analysis.

### A global peptide abundance correlation map

The SCC matrix of selected 21,363 peptides from V52 recorded how strongly or weakly each peptide is correlated with all other peptides. Although it is theoretically possible to be represented as a peptide interaction network with edges indicating strong correlations (Supplementary Figure 3). We visualized all 228,178,203 peptide-peptide correlations from 21,363 peptides as a global peptide correlation map using the t-SNE algorithm by embedding the peptides in a low-dimensional space (Figure 4A). In this map, the distance between peptides reflects the similarity in their abundance changes under various drug treatments. Notably, the global map was generated based on all pairwise peptide abundance changes under different perturbations in the metaproteomics datasets per individual, without any filtering on the SCC.

**Figure 4:**
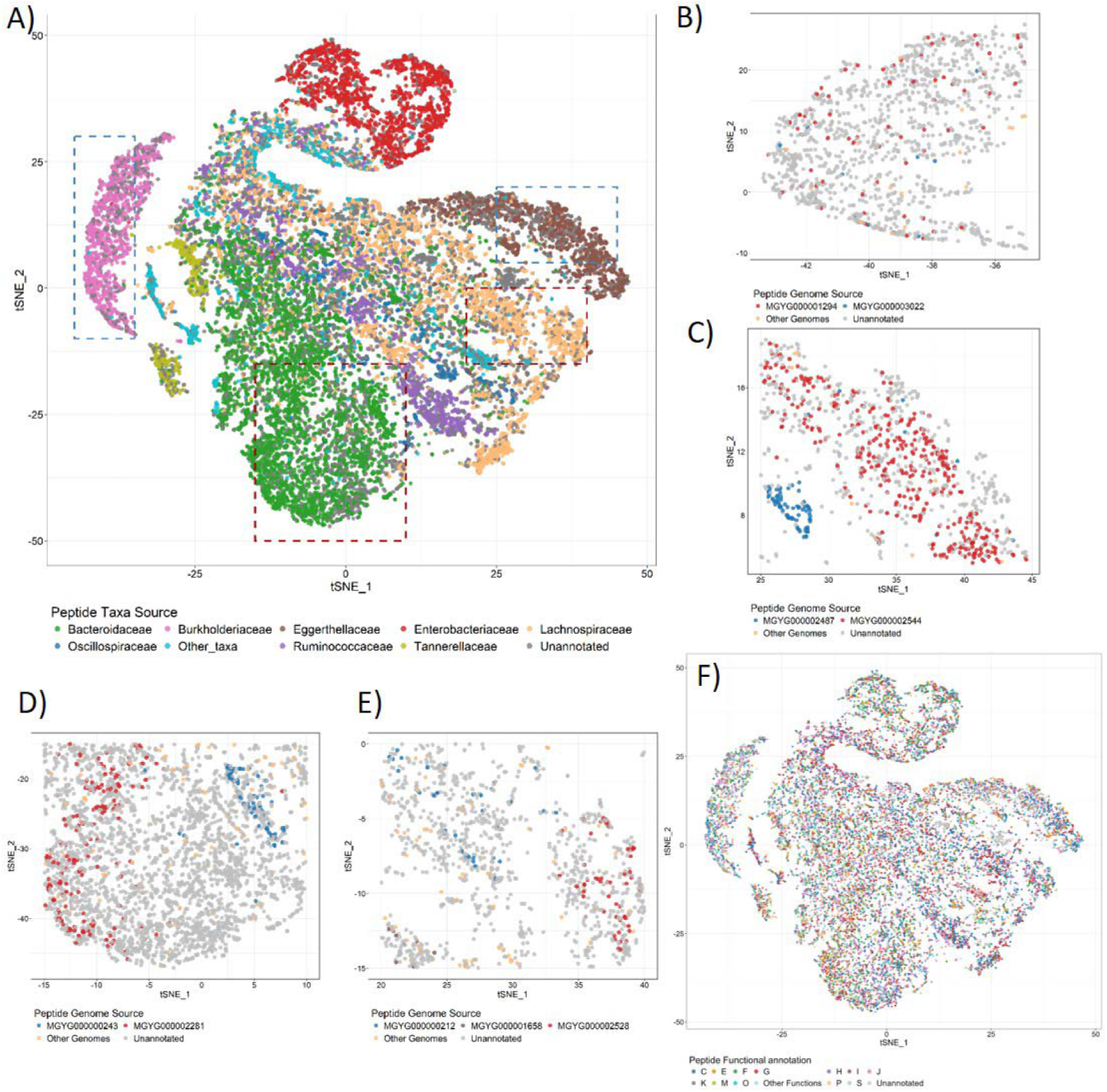
The global peptide correlation map of V52 generated using t-SNE. **A)** Peptide correlation map colored by family-level taxonomic annotations, showing the map broadly corresponds to taxa sources of peptides. The areas highlighted in panels B-E are boxed with dashed lines. **B-E)** Zoomed-in sections of the peptide correlation map for peptides from the different families: **B)** Burkholderiaceae, **C)** Eggerthellaceae, **D)** Bacteroidaceae, and **E)** Lachnospiraceae. Peptides are colored by genome-level taxonomic annotations. **F)** Peptide correlation map colored by functional annotations (COG categories), showing no overall clustering based on functional associations.

The peptide correlation map revealed a strong correlation between peptide abundance changes and their taxonomic sources. Peptides from the same taxon tend to cluster together in the correlation map across various taxonomic levels (Figure 4A and Supplementary Figure 4).

Specifically, at the family level, distinct clusters emerged, particularly for families such as Burkholderiaceae, Eggerthellaceae, and Enterobacteriaceae. Zooming into these families, most peptides with genome-level annotations were either assigned to a specific genome or grouped into sub-clusters corresponding to different genomes (Figures 4B and 4C). For other families, such as Bacteroidaceae and Lachnospiraceae, the cluster of peptides was less obvious. However, peptides were still grouped at the genome level in the zoomed-in maps (Figures 4D and 4E).

Although there is a clear clustering of peptides from the same taxon, the global peptide abundance correlation map showed less obvious clustering patterns based on protein function (Figure 4F and Supplementary Figure 5). However, when zooming into a single species, more distinct clusters emerged, representing peptides from proteins with similar or related functions (Supplementary Figure 6). This suggests that studying protein functional linkage through peptide abundance correlations may be more effective at the single-species level, as inter-species functional correlations are not easily discernible in the global map.

Overall, this global peptide correlation map provides an overall trend of peptide abundance correlations between different peptides and indicates that changes in taxa abundance are the primary driver of peptide abundance correlations across different perturbations.

### Peptide abundance correlations provide additional information on assigning peptide taxonomic source

Our results have shown that peptides from the same family clustered together in the global peptide correlation map (Figure 4A). However, within each family-level cluster, only a small proportion of peptides were annotated to specific genomes, while a larger proportion were only annotated at the family level, leaving their genome-level sources unclear (Figures 4B-4E and Supplementary Figure 4). Also as mentioned above, peptides from the same genome exhibit high SCCs in their abundance changes across different drug treatments (Figure 3C). Given a specific sample from the family Bacteroidaceae, genome-distinct peptides from the same genome were not grouped into a single cluster in the heatmap of the peptides SCC matrix. However, these peptides can still be grouped into several modules, each with high correlation coefficients (Figure 5A). Similar genome cluster modules were also observed in other families, such as Eggerthellaceae, Lachnospiraceae, and Burkholderiaceae (Supplementary Figure 7-9). This observation suggests that the SCC matrix can serve as a valuable input for machine learning models to assign peptide taxonomic sources.

**Figure 5:**
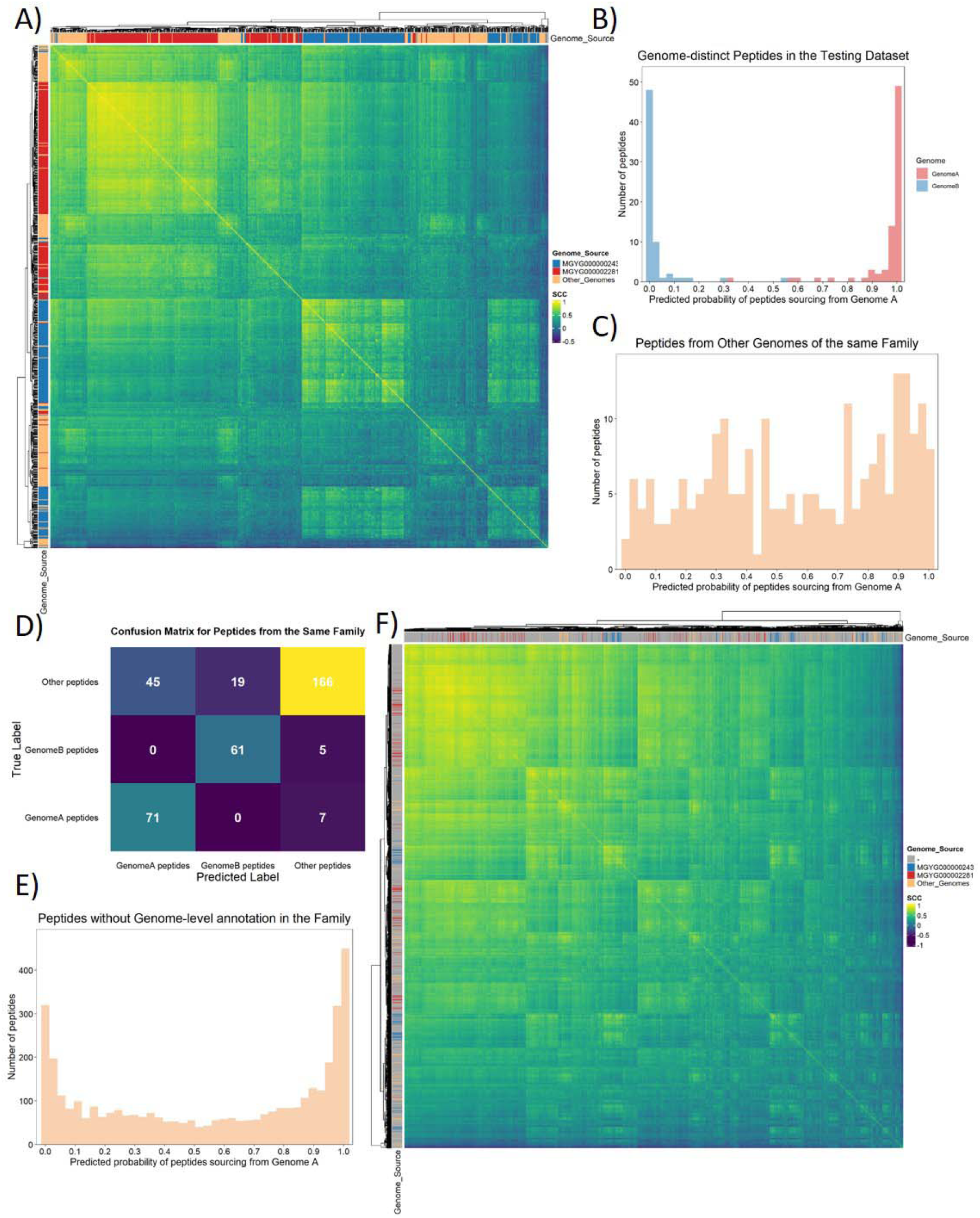
Applying peptide abundance correlations for peptide taxonomic assignments using the Bacteroidaceae family as an example. **A)** Heatmap of the SCC matrix for genome-distinct peptides from different genomes within the Bacteroidaceae family. **B)** Distribution of predicted probabilities for peptides sourcing from Genome A, focusing on genome-distinct peptides in the test dataset. **C)** Distribution of predicted probabilities for peptides sourcing from Genome A, focusing on peptides from other genomes within the same family. **D)** Confusion matrix from the Random Forest model applied to the combined dataset of genome-distinct peptides in the test dataset and peptides from other genomes within the family. **E)** Distribution of predicted probabilities for peptides sourcing from Genome A, focusing on genome-unannotated peptides within the Bacteroidaceae family. **F)** Heatmap of the SCC matrix for both genome-distinct peptides and genome-unannotated peptides within the Bacteroidaceae family. Genome-unannotated peptides were colored with grey in the color strip. Genome A, MGYG000002281; Genome B, MGYG000000243.

Using the trained Random Forest model for the family Bacteroidaceae, we found that most genome-distinct peptides from Genome A (MGYG000002281) and Genome B (MGYG000000243) in the test set were correctly classified with probabilities exceeding 90% (Figure 5B). Specifically, the model assigned these peptides to their respective genome sources with high confidence. In contrast, when applying the same 90% probability threshold to peptides from other genomes within the same family and peptides from other families, the model did not tend to attribute most of the peptides from other genomes within the same family (72.2%) and the peptides from other families (87.6%) to either Genome A or Genome B (Figure 5C and Supplementary Figure 10). The confusion matrices generated from the two test sets demonstrated that the trained model achieved high sensitivity (Figure 5D and Supplementary Figure 10). For peptides without genome-level annotations from the same family, the model classified a large amount of these peptides (48.9%) into either Genome A or Genome B (Figure 5E). This classification was further supported by the SCC heatmap, where unannotated peptides clustered closely with genome-distinct peptides from a specific genome (Figure 5F). In addition, models trained using the same method also effectively predicted the genome sources of peptides lacking genome-level annotations in other families, such as Eggerthellaceae, Lachnospiraceae, and Burkholderiaceae (Supplementary Figures 7-9).

### Functionally related peptides were connected in single-species peptide abundance correlation networks

Our results revealed limited abundance correlations between peptides from proteins with the same or related functions (Figures 3E-3F and Supplementary Figure 5). Additionally, we found that taxa abundance changes are the major driver of peptide abundance changes across different perturbations or drug treatments. Consequently, peptides from proteins with the same or related functions were less likely to be correlated across different taxa compared to peptides originating from the same taxon. Therefore, it is more appropriate to analyze peptide functional linkages at the single-species level by focusing on single-species subsets of the metaproteomics dataset.

The strong effect of taxonomic abundance changes could cause peptides from the same taxon to show high correlations, even if they are involved in different functions (Supplementary Figure 11). To minimize this effect and better study the correlations between functionally related peptides, we calculated taxon-based normalized peptide abundance (TNPA) from original peptide abundance (OPA) for the top 10 species/genomes with the highest number of identified peptides, as described in the methods section. TNPA was mostly moderately or weakly positively correlated with OPA in most studied species (Supplementary Figure 12), and we observed that some peptides with high SCCs for OPA log2-FC exhibited low SCCs for TNPA log2-FC (Figure 6A and Supplementary Figure 13). This suggests that many peptide abundance correlations driven by taxa abundance changes were not significantly correlating at the TNPA level as shown in Supplementary Figure 10.

**Figure 6:**
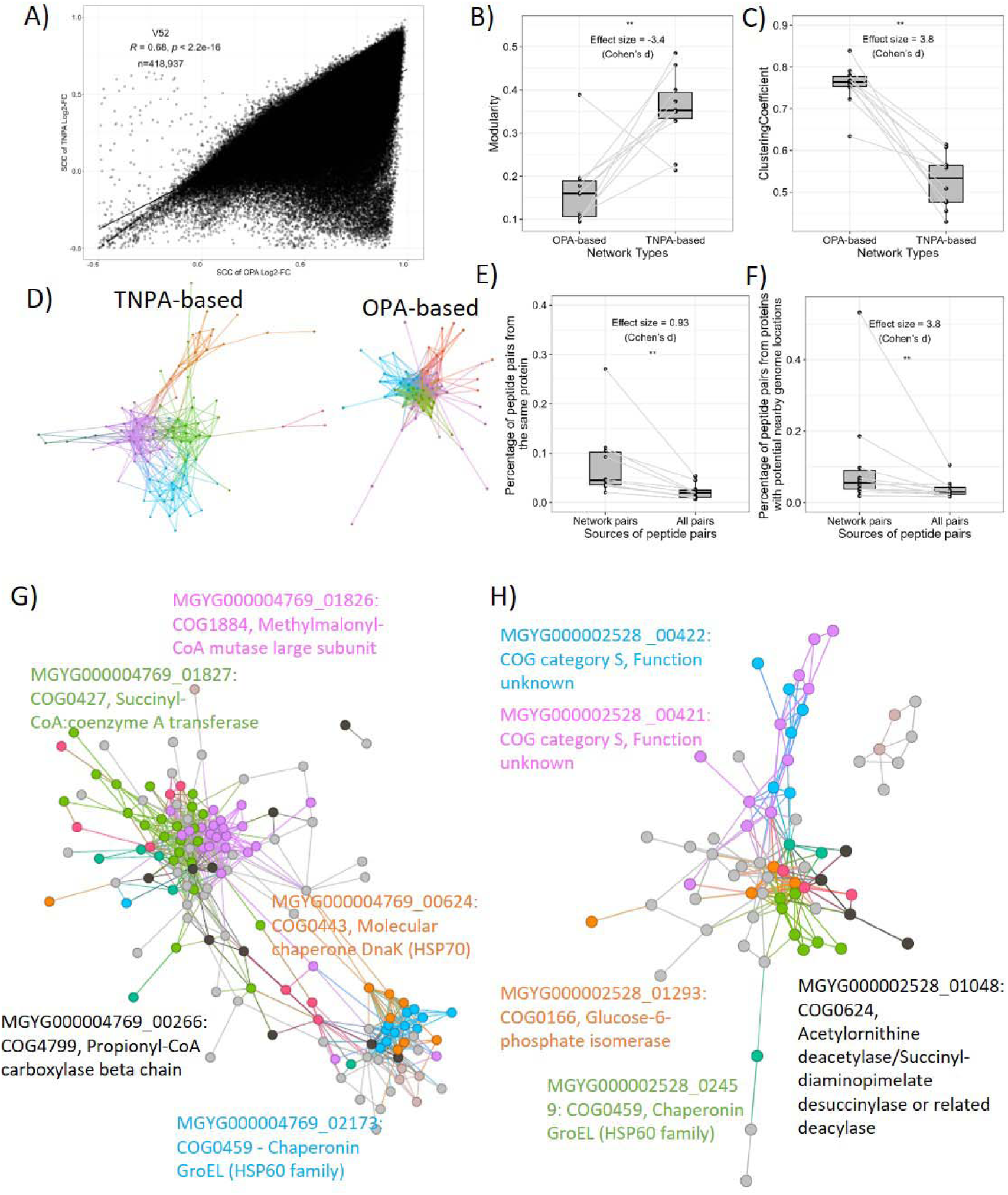
Functionally related peptides were connected in V52 single-species peptide abundance correlation networks constructed with SCCs of TNPA log2-FC. **A)** Correlation between SCCs of peptide log2-FC based on TNPA and SCCs based on OPA (original peptide abundance) in the top ten genomes from V52. **B)-C)** Higher modularity and lower clustering coefficient of peptide correlation networks constructed with SCC of TNPA log2-FC than networks constructed with SCC of OPA log2-FC. **D)** Comparison of representative single-species peptide correlation networks constructed with TNPA and OPA. Nodes are colored according to their modularity class from Gephi. **E)** Proportion of peptide pairs from the same protein in single-species peptide correlation networks compared to the proportion in all peptide pairs from the corresponding species. **F)** Proportion of peptide pairs from proteins potentially located near each other in the genome (≤ 10 gene ID difference) in single-species peptide correlation networks compared to the proportion in all peptide pairs from the corresponding species. ** indicates statistical significance at the p ≤ 0.01 levels by two-sided Mann-Whitney U test. **G)-H)** Single-species peptide abundance correlation networks for MGYG000004769 and MGYG000002528. Peptides from proteins with only one peptide in the network were removed. Peptides from the top eight proteins with the most peptides in each network are annotated with distinct colors, and peptides from other proteins are colored grey.

To reveal functional linkages, single-species peptide abundance correlation networks were constructed using peptide pairs with the top 5% SCC from each of the top 10 species with the most identified peptides. Two sets of networks were constructed using SCCs derived from OPA and TNPA, respectively (Supplementary Figures 14 and 15). Networks constructed with TNPA had higher modularity (Figure 6B) and lower clustering coefficients (Figure 6C), indicating that these networks had dense connections within modules but sparse connections between different modules. The lower clustering coefficients also suggested that TNPA-based networks were less tightly connected as shown in the Figure 6D.

Single-species peptide abundance correlation networks constructed with TNPA revealed functional linkages. Peptides from the same protein and those from proteins potentially located near each other in the genomes (with ≤ 10 gene ID differences in the UHGG) were considered functionally related. Peptide pairs in the network showed a higher percentage of these types of relationships compared to all peptide pairs (Figure 6E and Figure 6F). It is worth noting that gene ID differences do not always perfectly reflect physical genome distance; however, they generally indicate proximity when genes are located on the same contig. Here, we provide examples of connections between peptides from functionally related proteins. In the peptide correlation network of genome MGYG000004769 (*Phascolarctobacterium faecium*), peptides from MGYG000004769_01826 (Methylmalonyl-CoA mutase large subunit) and MGYG000004769_01827 (Succinyl-CoA: coenzyme A transferase) were densely connected (Figure 6G). Genes encoding these two proteins are located near each other in the genome, with adjacent gene ID numbers and only a 218bp intergenic region on the same contig. These proteins also have closely related functions in the TCA cycle. Methylmalonyl-CoA mutase catalyzes the conversion of methylmalonyl-CoA to succinyl-CoA, while Succinyl-CoA:coenzyme A transferase then catalyzes the reversible reaction: succinyl-CoA + L-malate ⇄ succinate + L-malyl-CoA. In the same network, peptides from MGYG000004769_00624 (Molecular chaperone DnaK) and MGYG000004769_02173 (Chaperonin GroEL), which have similar functional annotations, were also connected. In another peptide correlation network of genome MGYG000002528 (*Anaerostipes hadrus*), peptides from proteins MGYG000002528_01044 (NAD-dependent dihydropyrimidine dehydrogenase subunit PreA) and MGYG000002528_01048 (Allantoate amidohydrolase) were densely connected (Figure 6H). These proteins are also encoded by genes with adjacent gene numbers which are separated by a 2,717 bp intergenic region.

In summary, single-species peptide abundance correlation networks constructed using TNPA revealed connections between functionally related peptides. These connections demonstrated the potential for using peptide abundance correlations to predict protein functions.

### Applying single-species peptide correlation networks for predicting unknown protein functions

An average of 5.2% of peptides in the 10 constructed single-species peptide correlation networks originated from proteins of unknown functions (PUFs). This percentage was comparable to the overall proportion of peptides from unannotated proteins in each species (Figure 7A). To predict functions for PUFs we calculated peptide abundance correlations between peptides from PUFs and peptides from annotated proteins. The proportion of connections between peptides from PUFs and peptides from annotated proteins varied across species (Figure 7B). For genome MGYG000002478 (*Phocaeicola dorei*), the proportion reached 24%, while in other genomes, the lowest was 2%. Here, we provide examples from the top two genomes with the highest percentage of connections between peptides from PUFs and peptides from annotated proteins to show the feasibility of predicting microbial protein functions using peptide abundance correlations.

**Figure 7:**
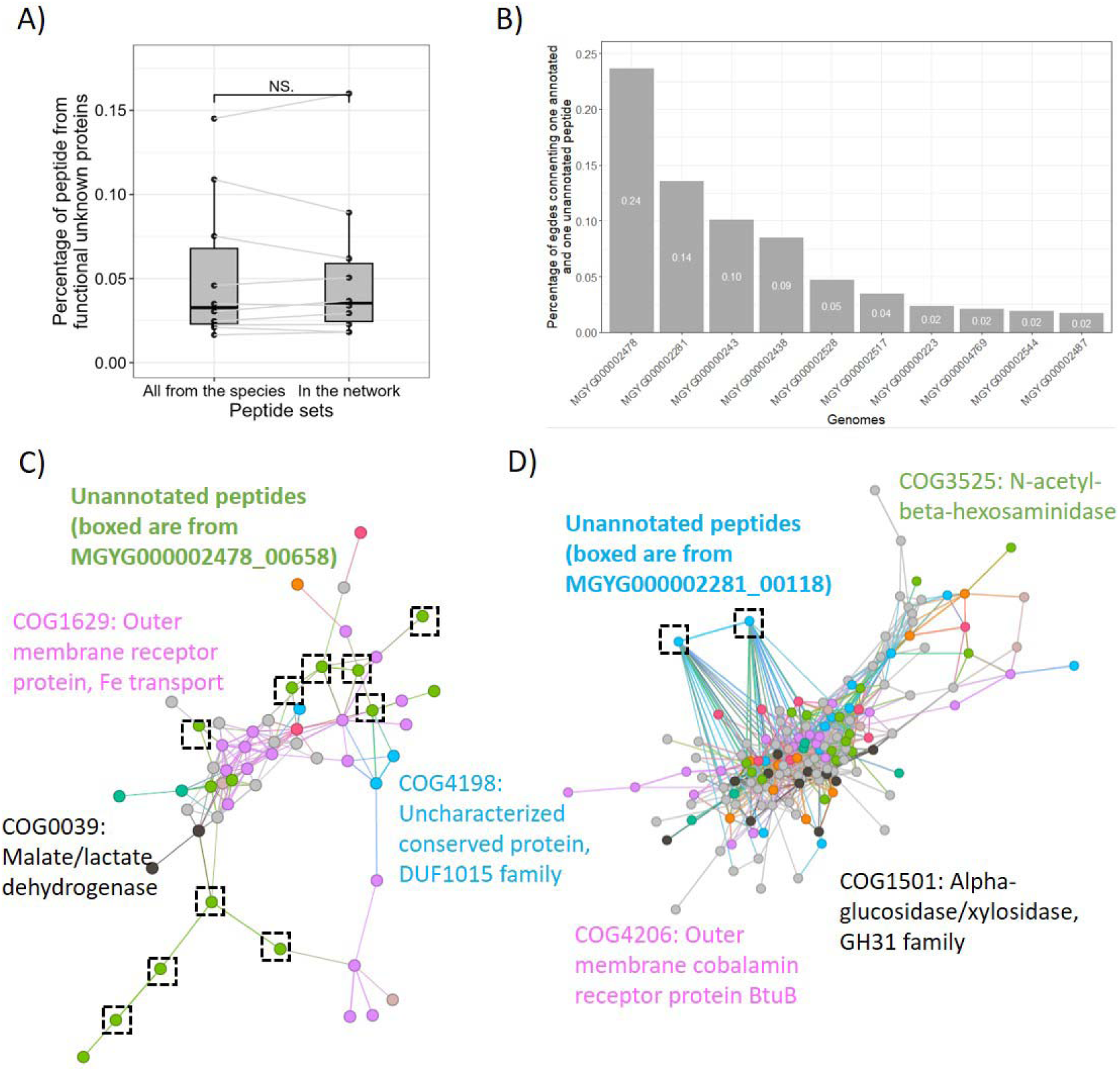
Predicting functions of proteins of unknown functions (PUFs). **A)** Proportion of peptides from PUFs in each single species peptide correlation network and proportion of peptides from PUFs in all identified peptides from the corresponding species. NS. indicates statistical significance at the p > 0.05 levels by two-sided Mann-Whitney U test. **B)** Distribution of the proportions of edges between peptides from PUFs and peptides from proteins with known function in the peptide correlation network of each species. **C)-D)** Peptide abundance correlation networks for the two species, MGYG000002478 and MGYG000002281, with the highest percentage of edges between peptides from PUFs and peptides from proteins with known function. Networks were constructed based on peptide pairs with the top 5% SCCs from each species. Peptides from proteins with only one peptide in the network were removed. Peptides from the top eight COG families in each network are annotated with distinct colors, and peptides from other COG families are colored grey.

In the network of genome MGYG000002478 (Figure 7C), *Phocaeicola dorei*, peptides from uncharacterized protein MGYG000002478_00658 had the largest number of connections (31) to annotated peptides in the network. Of these, 10 connections were to peptides annotated to COG1629 (Outer membrane receptor protein, Fe transport), with 7 linked to peptides from MGYG000002478_00657, an iron complex outer-membrane receptor protein (K02014). This indicates that the uncharacterized protein is likely to relate to tran-membrane transport. This prediction was further supported by the InterProScan of the uncharacterized protein, showing it has a SusD-like domain, a typical outer membrane protein feature. In addition, most parts of the protein were assigned to a non-cytoplasmic domain, also indicating its extracellular location.

In the network of genome MGYG000002281 (Figure 7D), *Bacteroides faecis*, peptides from uncharacterized protein MGYG000002281_00118 had the largest number of connections (52) to annotated peptides in the network. Excluding 9 peptide connections to functionally ambiguous COG0457 (Tetratricopeptide repeat protein), the second-largest group comprises 7 connections to COG3525 (N-acetyl-beta-hexosaminidase), including 3 connections to MGYG000002281_02589 (hyaluronoglucosaminidase, K01197), 2 to MGYG000002281_04635( hexosaminidase, K12373), and 2 to MGYG000002281_01369 (hexosaminidase, K12373). These connections suggest the uncharacterized protein is likely to be involved in carbohydrate metabolism. An InterProScan analysis of the protein revealed a non-cytoplasmic domain, indicating that it is a membrane-bound protein predicted to be outside the membrane, possibly acting as a signal protein for carbohydrate metabolism. In contrast, searching this protein in the AlphaFold Protein Structure Database^37,38^ matched UniProt ID A0A1E9BZD4 (100% identity, HSP score of 1197), an uncharacterized protein without known biological function. These findings highlight the value of peptide correlation networks in providing complementary insights for predicting the potential biological role of PUFs.

## Discussion

We analyzed peptide abundance correlations in a metaproteomics dataset of *in vitro* cultured human gut microbiomes with different drug treatments/perturbations, demonstrating the feasibility of applying peptide abundance correlations for peptide taxonomic assignments as well as revealing protein functional linkages in single-species subsets.

Protein abundance correlations have been applied to study protein functional linkage and to predict functions of functional unknown proteins in single-species proteomics datasets ^13–15^. In these single-species studies, functionally associated proteins have coordinated changes in abundance across perturbations ^13–15^, and proteins with identical subcellular localization also exhibited coordinated abundance changes ^13^. However, in our metaproteomics dataset, the abundances of peptides from proteins of the same taxon instead of peptides from proteins with associated functions were more likely to have correlated abundance changes.

Peptides from the same taxon showing high abundance correlation are largely due to the significant impact of environmental disturbances on the composition of microbial communities^39^. In contrast, the limited correlation of peptides from proteins with related functions can be attributed to the high functional redundancy of human gut microbial communities^40^. Specifically, a decrease in peptides from proteins with a particular function could lead to an increase in peptides from proteins performing a similar function in a phylogenetically unrelated species. This compensatory mechanism helps maintain the overall functional stability of the microbial community, resulting in limited correlations among peptides from proteins with related functions.

Our study is the first to investigate peptide abundance correlations in a gut microbiome metaproteomic dataset. However, similar studies have been conducted using metagenomic data, such as genetic correlation networks from soil metagenomes which have revealed a hierarchical functional structure^16^, similar to findings in single-species cellular genetic correlation networks^41^. In contrast, our analysis of peptide abundance correlations primarily showed a clear clustering of peptides from the same taxon, with a less obvious functional hierarchy. This underscores that functional insights derived from protein abundance measurements are different from those inferred from genetic materials through metagenomics or metatranscriptomics^42^.

Our global peptide abundance correlation map also provided clues for studying microbial responses to drugs. In the constructed peptide correlation maps (Figure 2A), peptides from the family Enterobacteriaceae were usually separated from other taxa, indicating unique response patterns for this taxonomic group. This is supported by previous experimental studies. For example, psychotropic drugs like fluoxetine have shown antimicrobial activity against *Escherichia coli* (family Enterobacteriaceae), while the abundance of other human gut species increased in the *in vitro* microbiome after the treatment of the same drug ^43^. In addition, microbial genera have been classified into three distinct taxonomic clusters, each representing a different pattern of drug response^26^. Patterns resembling these taxonomic clusters can also be observed in our peptide correlation maps. For instance, peptides from *Parasutterella* (Burkholderiaceae family) and *Eggerthella* (Eggerthellaceae family), which belong to two clusters exhibiting distinct response patterns, are also well separated in our peptide correlation map (Figure 2A and Supplementary Figure 4).

Higher correlations of peptides from the same taxon also provide a solution for peptide taxonomic source assignment which remains a challenge for peptide-centric metaproteomics analysis. Although most of the in-silico digested peptides were genome-distinct peptides ^44^, most of the peptides identified from real metaproteomics datasets were shared by different genomes ^21^. In this study, even after refining the potential genome sources, a large number of peptides could only be assigned to a family-level LCA, and the actual taxonomic sources of these peptides remain unknown (Figures 4B-4E). By investigating peptide abundance correlations, additional information was provided for peptide taxonomic source assignment. This is expected to provide a microbial biomass profile with higher accuracy using metaproteomics data ^45^.

Although taxa-abundance changes were major contributors to peptide abundance correlations, to reveal peptide functional linkages, peptide abundance correlations were further analyzed at the single-species level by extracting subsets of genome-distinct peptides of each genome from the metaproteomics dataset. In addition, to reduce the impact of taxa-abundance changes, TNPA was calculated. The principle of the calculation of TNPA was similar to a normalization method named LFQRatio ^46^ which divided protein LFQ intensity by the sum of all protein LFQ intensities for its respective strain in a microbial coculture system. LFQRatio has shown its ability to transform absolute protein quantification data into accurate and biologically meaningful protein abundance values for samples with multiple species at variable cell ratios.

Similarly, in our study, we also found that taxa-based normalized peptide abundance (TNPA) showed a good ability to reveal functional linkages (Figures 6E and 6F). Peptides from proteins with related functions were connected in the single-species peptide abundance correlation network constructed with TNPA.

A limitation of this study comes partly from metaproteomics itself. Current metaproteomics has limited proteome coverage ^47,48^. It is estimated that around 200 bacterial species reside in the human gut microbial community ^49^. However, bacterial species with an abundance lower than 0.5% were hard to detect with current metaproteomics techniques ^50^, and the number of peptides and proteins identified in low-abundant species is limited ^48^. To that extent, only the top ten genomes were investigated to study intra-species functional relations in this work. Even in these genomes, the number of peptides was still limited. This results in the limited scale of constructed single-species peptide correlation networks. A lot of peptides from proteins with interesting or unknown functions were not identified and were not included in the network. In addition to considering proteome coverage, it is also worth noting that, a high-resolution LC-MS/MS for identification and quantification is also needed. Our analysis in another metaproteomics dataset from Q Exactive mass spectrometer with the same procedure showed a weak abundance correlation of same-taxon peptides (Supplementary Figure 16). Fortunately, DIA-based metaproteomics has proven to be able to significantly increase the number of identified peptides as well as to improve quantification reproducibilities ^51,52^ which has the potential to expand and strengthen applying peptide abundance correlations to reveal more interesting findings.

Another aspect that needs improvement is distinguishing true functional correlations from incidental ones in single-species peptide correlation networks. First, the number of samples used to calculate Spearman correlation coefficients should be carefully considered, as these coefficients require sufficient sample sizes to achieve high confidence in the correlations detected ^53^. Additionally, applying a strict threshold or employing more refined methods to filter out functionally related interactions from correlations driven by other factors is crucial, since abundance correlations do not always indicate functional relationships ^54^. Ultimately, the field would benefit from larger, more reproducible datasets and refined protein association reference methods to advance the study of peptide/protein abundance correlations.

## Conclusion

In summary, fluctuations in peptide abundance contain wealthy information on the microbiome’s response to perturbations. These fluctuations can be harnessed to assign peptides to their protein and taxon of origin as well as to predict function for functional unknown proteins. In the metaproteomics dataset, peptides from the same taxon were clustered in the peptide correlation map, suggesting that microbiome taxonomic abundance change is the major contributor to peptide abundance changes. We anticipate that the concept of peptide/protein abundance correlations needs further investigation to deepen taxonomic and functional understanding of metaproteomics data.

## Author Contributions

CRediT authorship contribution statement. Zhongzhi Sun: Data curation, Formal analysis, Investigation, Methodology, Visualization, Writing – original draft. Zhibin Ning: Conceptualization, Methodology, Investigation, Writing – original draft. Qing Wu: Methodology, Investigation, Writing – review & editing. Andrew Doxey: Methodology, Investigation, Writing – review & editing. Daniel Figeys: Conceptualization, Funding acquisition, Supervision, Writing – original draft.

## Acknowledgments

Substantial financial support was provided by the Natural Sciences and Engineering Research Council of Canada (NSERC) through the discovery grant (to D. F.). Z. S. and Q. W. were funded by a stipend from the NSERC CREATE in Technologies for Microbiome Science and Engineering (TECHNOMISE) Program.

