## Supplementary Figure for "Peptide abundance correlations in metaproteomics enhance taxonomic and functional analysis of the human gut microbiome"

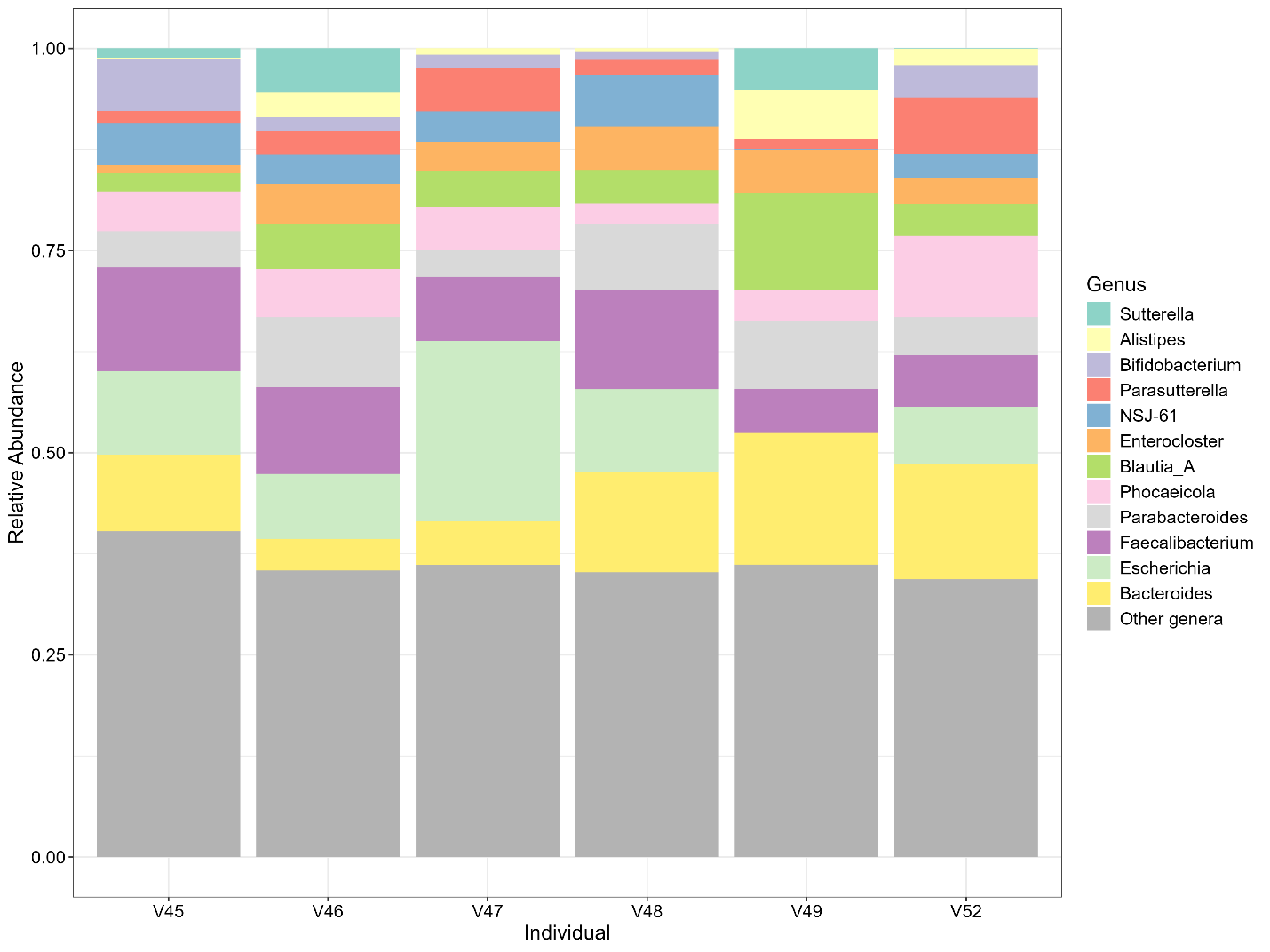

### **Supplementary Figure 1.**

**The relative abundance of peptides assigned to different genera across individuals.** Only peptides with genus-level taxonomic annotations were included in the plot. The top 12 genera, ranked by the total number of peptides across all individuals, are shown in distinct colors, while peptides assigned to other genera are grouped under the "Other genera" category.

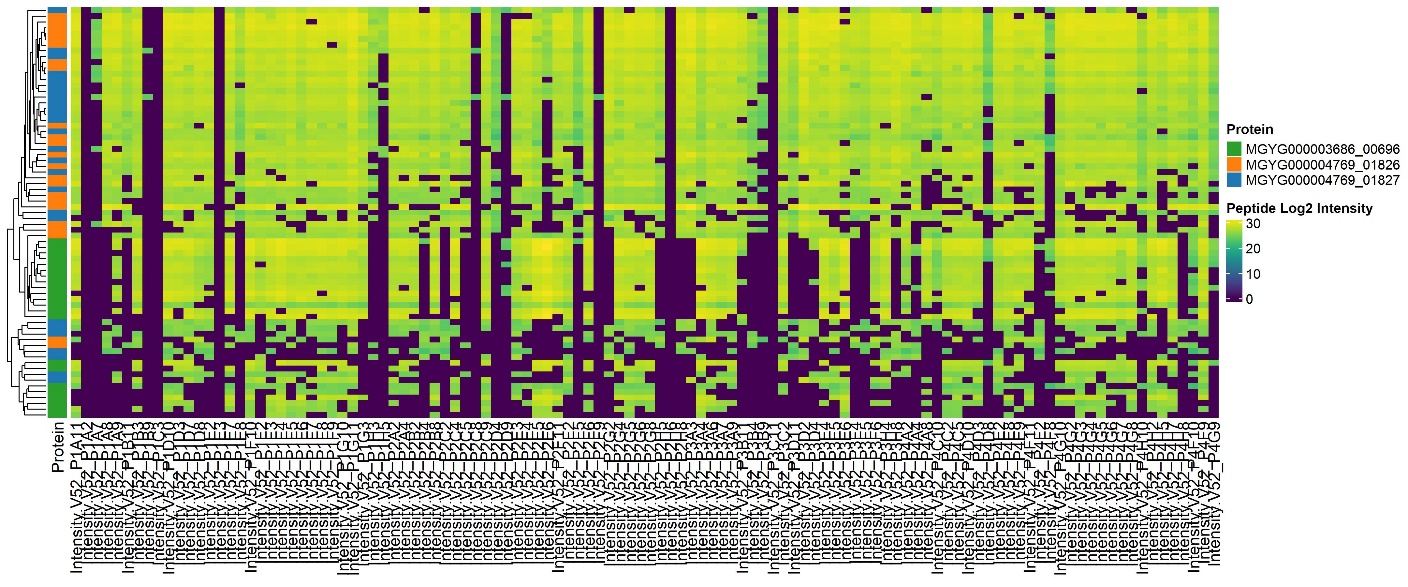

### **Supplementary Figure 2.**

**Peptide abundance profiles from three distinct proteins.** Peptides from the same proteins showed similar abundance profiles across samples under different treatments/perturbations.

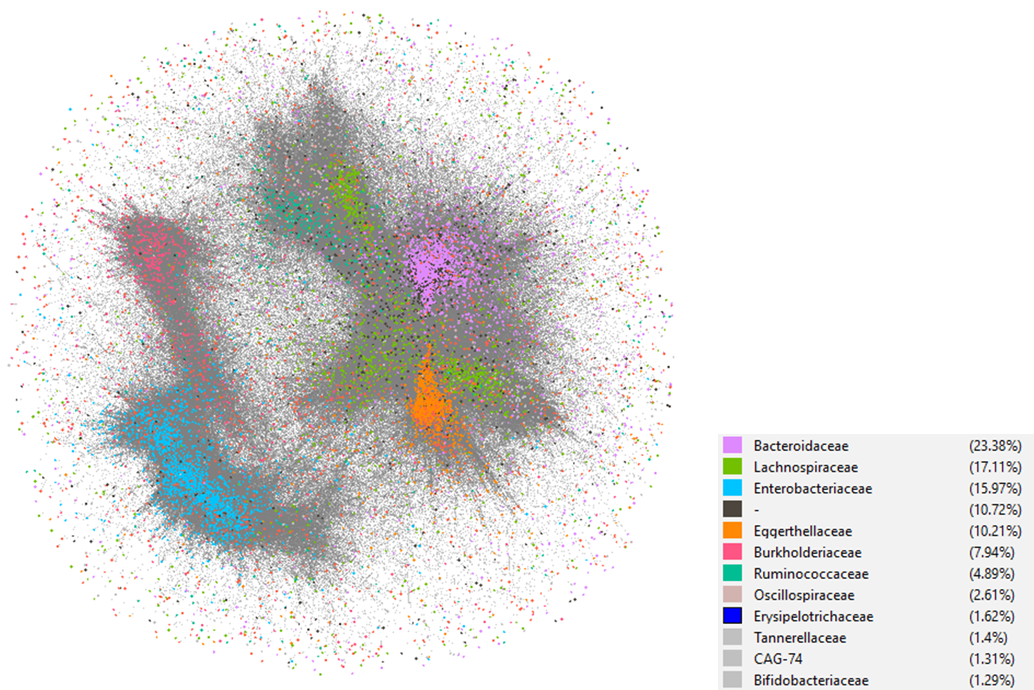

### **Supplementary Figure 3.**

**Global peptide abundance correlation network of V52.** The network comprises peptide pairs with the top 1% highest Spearman correlation coefficients (SCC). It consists of 12,549 nodes (representing individual peptides) and 2,281,783 edges (representing peptide pairs with high SCC).

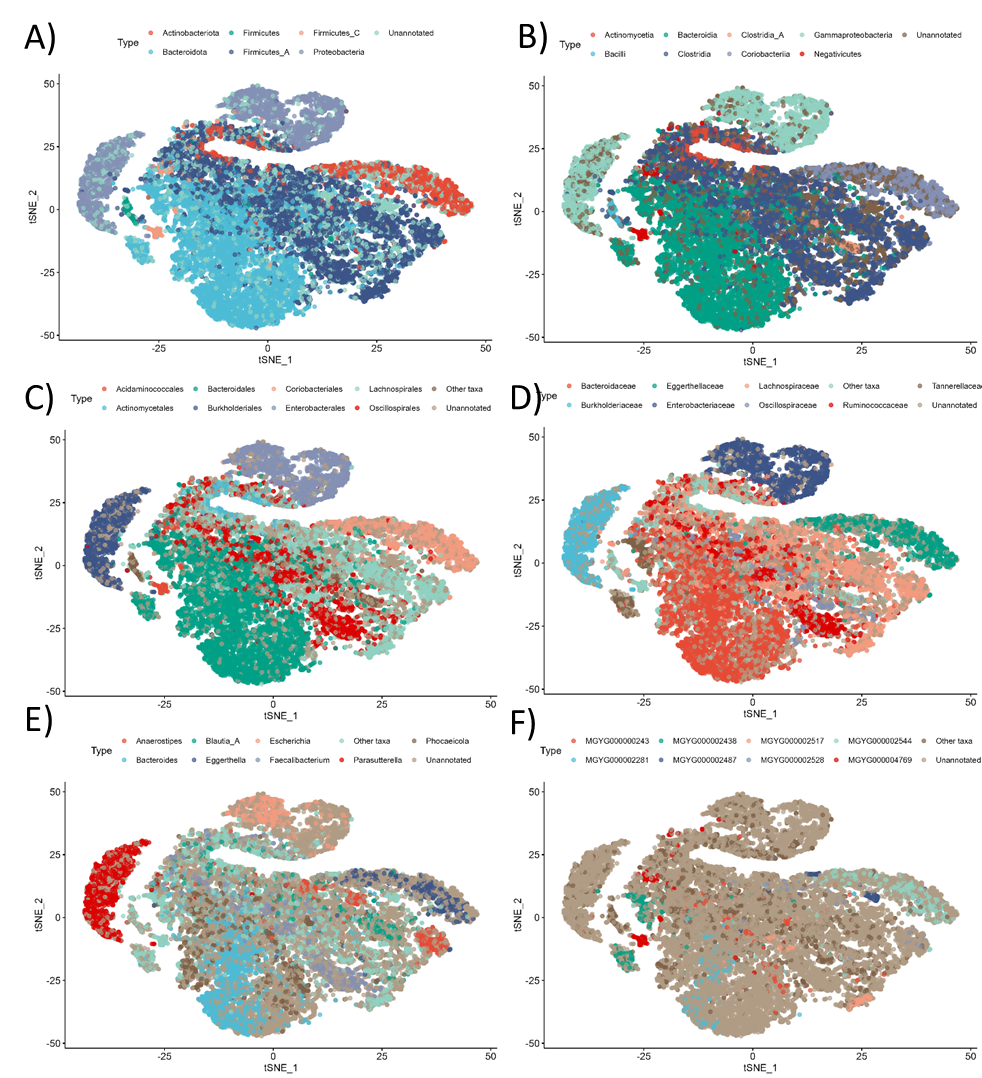

### Supplementary Figure 4.

**Global peptide correlation maps of V52 colored at different taxa levels.** A) Phylum, B) Class, C) Order, D) Family, E) Genus, and F) Species (Genome).

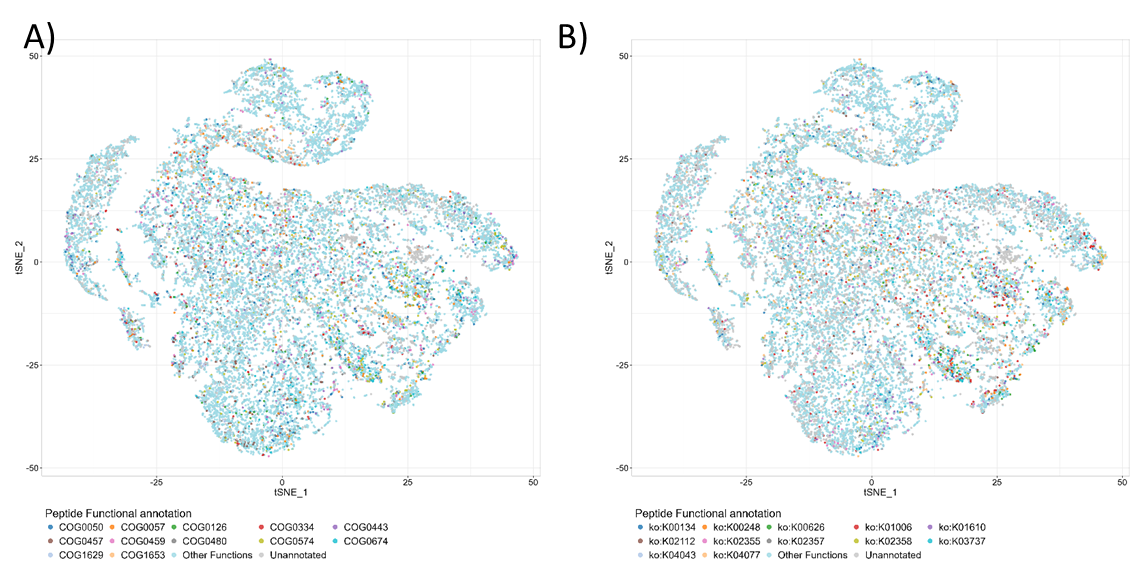

### Supplementary Figure 5.

**Global peptide abundance correlation maps of V52 colored at different functional categories.** A) COG family and B) KEGG ko.

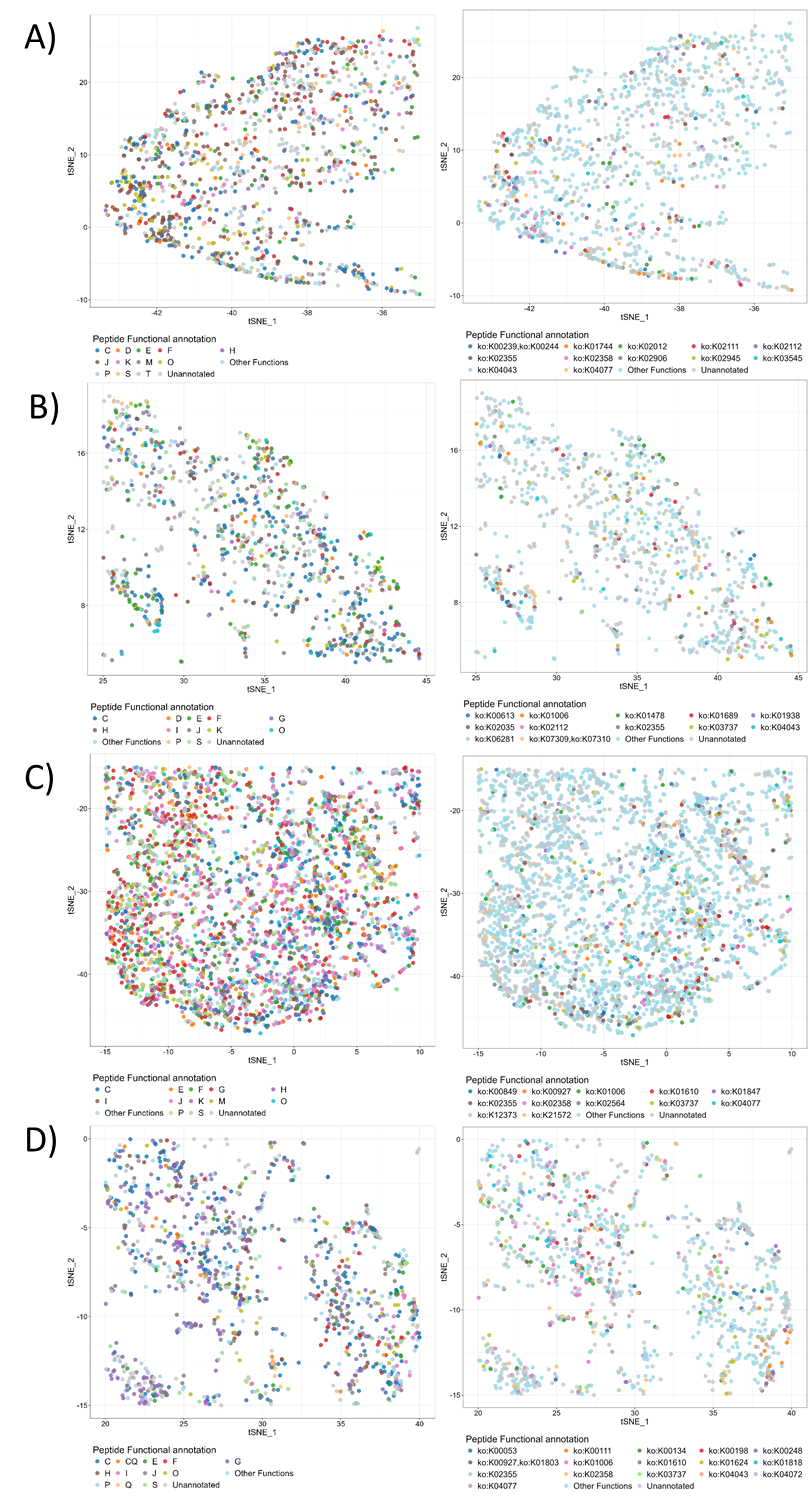

### Supplementary Figure 6.

**Zoomed-in peptide abundance correlation maps of V52 colored at functional categories: COG family (left) and KEGG ko (right).** Zoomed in maps for bacteria families: A) Burkholderiaceae, B) Eggerthellaceae, C) Bacteroidaceae, and D) Lachnospiraceae.

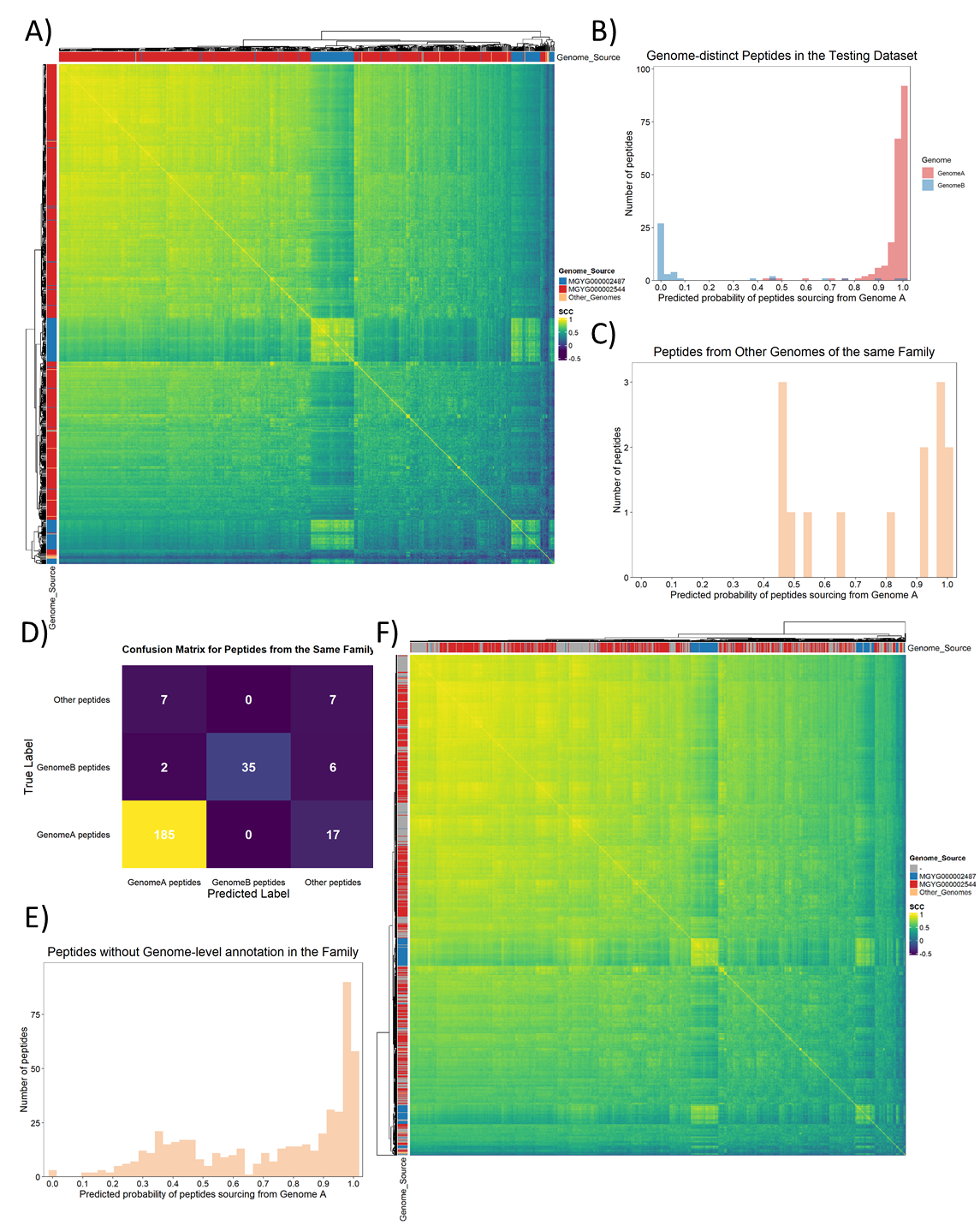

### Supplementary Figure 7.

**Applying peptide abundance correlation for peptide taxonomic assignments in the family Eggerthellaceae.** A) Heatmap of the SCC matrix for genome-distinct peptides from different genomes within the family. B) Distribution of predicted probabilities for peptides sourcing from Genome A, focusing on genome-distinct peptides in the test dataset. C) Distribution of predicted probabilities for peptides sourcing from Genome A, focusing on peptides from other genomes within the same family. D) Confusion matrix from the Random Forest model applied to the combined dataset of genome-distinct peptides in the test dataset and peptides from other genomes within the family. E) Distribution of predicted probabilities for peptides sourcing from Genome A, focusing on genome-unannotated peptides within the family. F) Heatmap of the SCC matrix for both genome-distinct peptides and genome-unannotated peptides within the family. Genome A, MGYG000002544; Genome B, MGYG000002487.

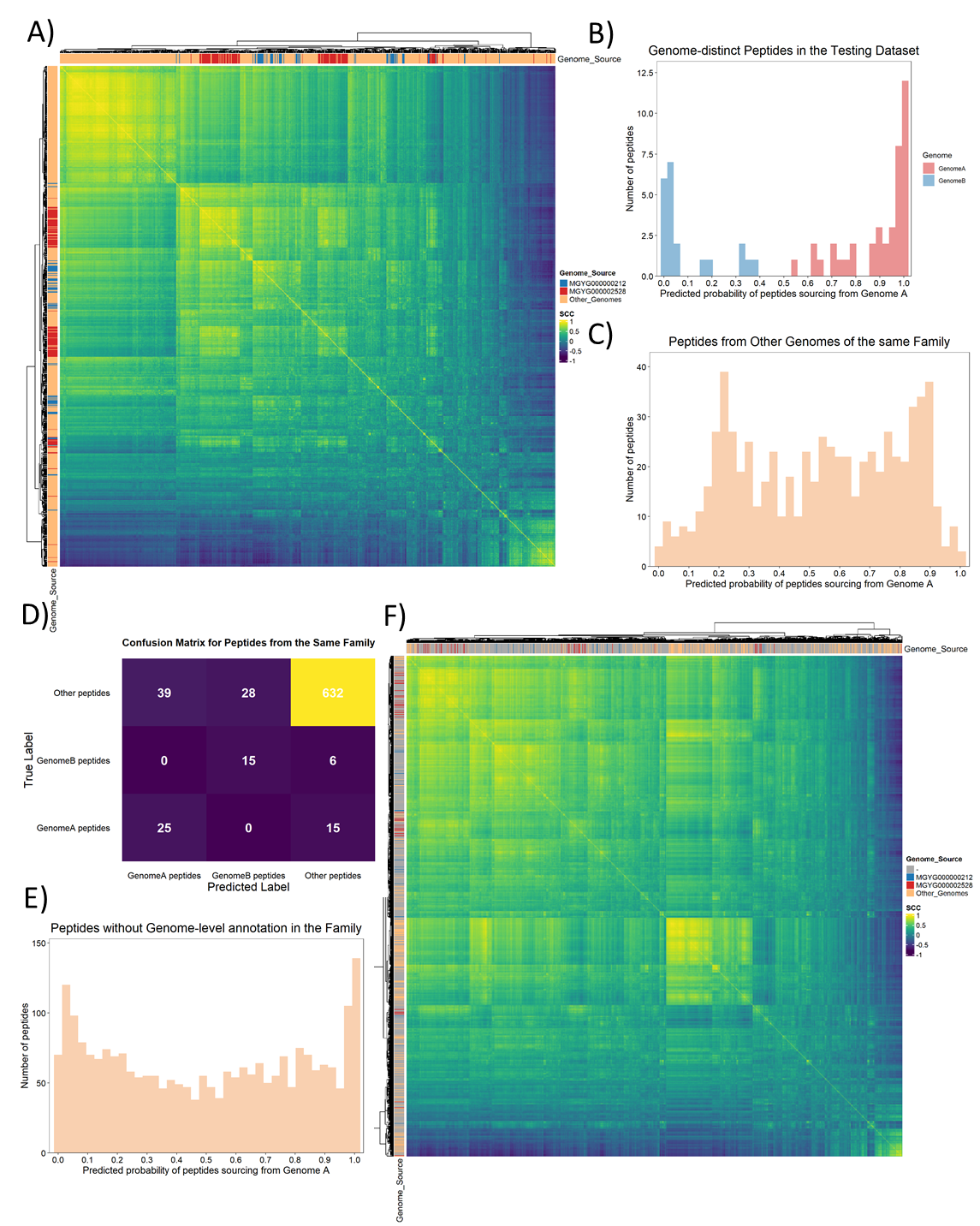

### Supplementary Figure 8.

**Applying peptide abundance correlation for peptide taxonomic assignments in the family Lachnospiraceae.** A) Heatmap of the SCC matrix for genome-distinct peptides from different genomes within the family. B) Distribution of predicted probabilities for peptides sourcing from Genome A, focusing on genome-distinct peptides in the test dataset. C) Distribution of predicted probabilities for peptides sourcing from Genome A, focusing on peptides from other genomes within the same family. D) Confusion matrix from the Random Forest model applied to the combined dataset of genome-distinct peptides in the test dataset and peptides from other genomes within the family. E) Distribution of predicted probabilities for peptides sourcing from Genome A, focusing on genome-unannotated peptides within the family. F) Heatmap of the SCC matrix for both genome-distinct peptides and genome-unannotated peptides within the family. Genome A, MGYG000002528; Genome B, MGYG000000212.

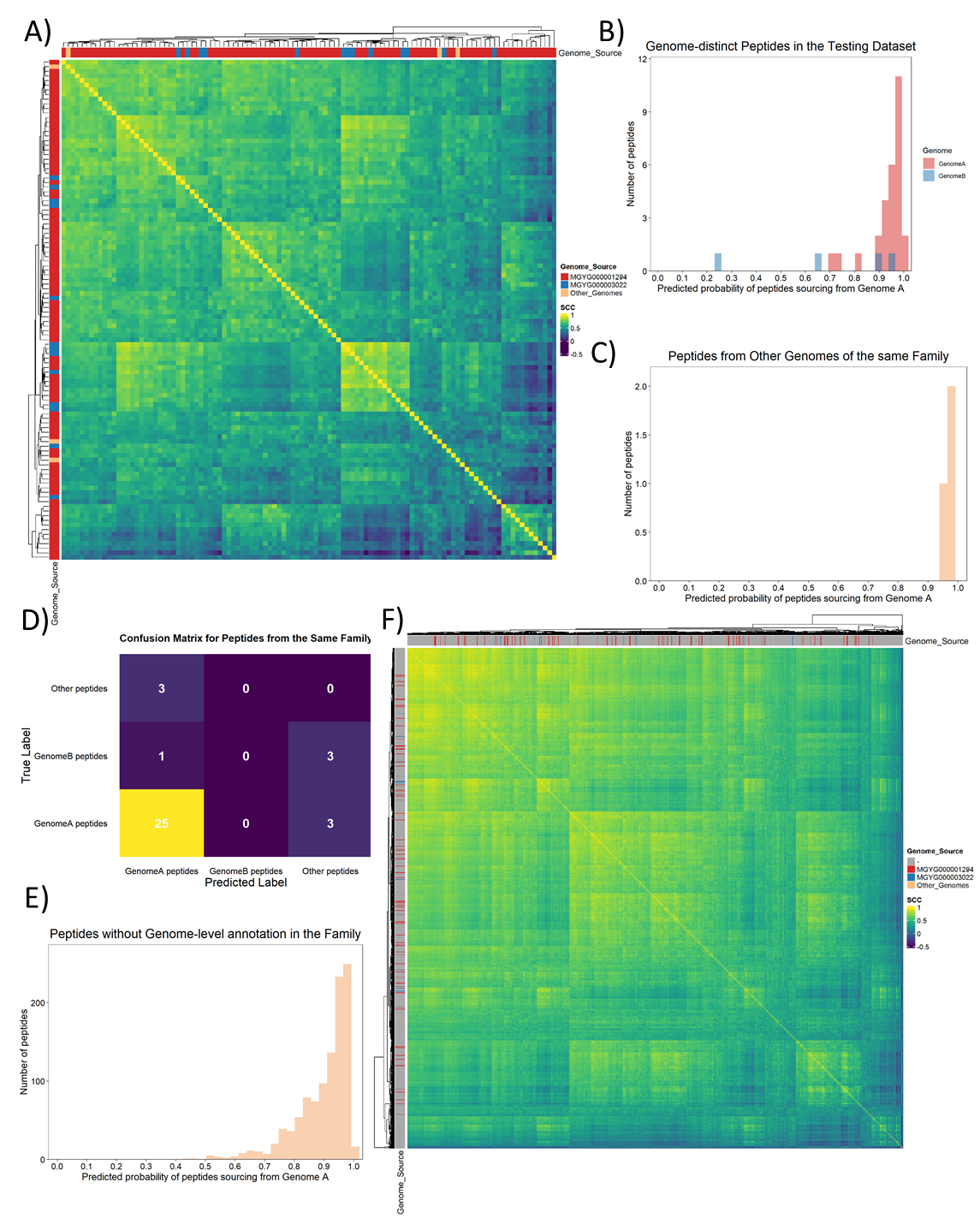

### Supplementary Figure 9.

**Applying peptide abundance correlation for peptide taxonomic assignments in the family Burkholderiaceae.** A) Heatmap of the SCC matrix for genome-distinct peptides from different genomes within the family. B) Distribution of predicted probabilities for peptides sourcing from Genome A, focusing on genome-distinct peptides in the test dataset. C) Distribution of predicted probabilities for peptides sourcing from Genome A, focusing on peptides from other genomes within the same family. D) Confusion matrix from the Random Forest model applied to the combined dataset of genome-distinct peptides in the test dataset and peptides from other genomes within the family (Genome B, MGYG000003022). E) Distribution of predicted probabilities for peptides sourcing from Genome A, focusing on genome-unannotated peptides within the family. F) Heatmap of the SCC matrix for both genome-distinct peptides and genome-unannotated peptides within the family. Genome A, MGYG000001294; Genome B, MGYG000003022.

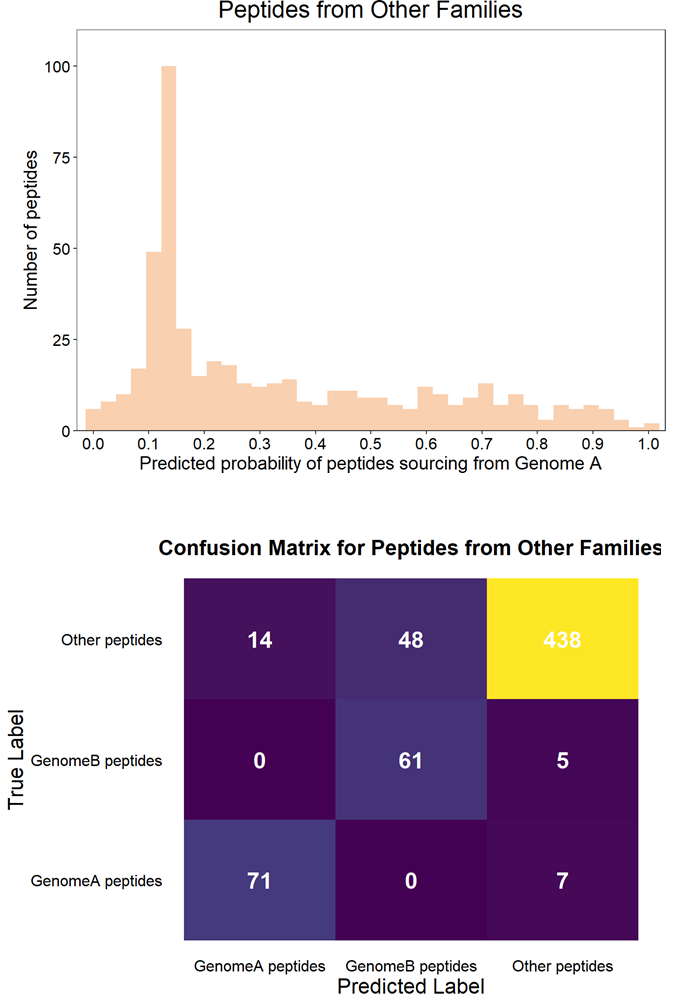

### Supplementary Figure 10.

**Peptide taxonomic source prediction results for testing peptides from families other than Bacteroidaceae.** The prediction model was trained using genome-distinct peptides from Bacteroidaceae. Genome A, MGYG000002281; Genome B, MGYG000000243.

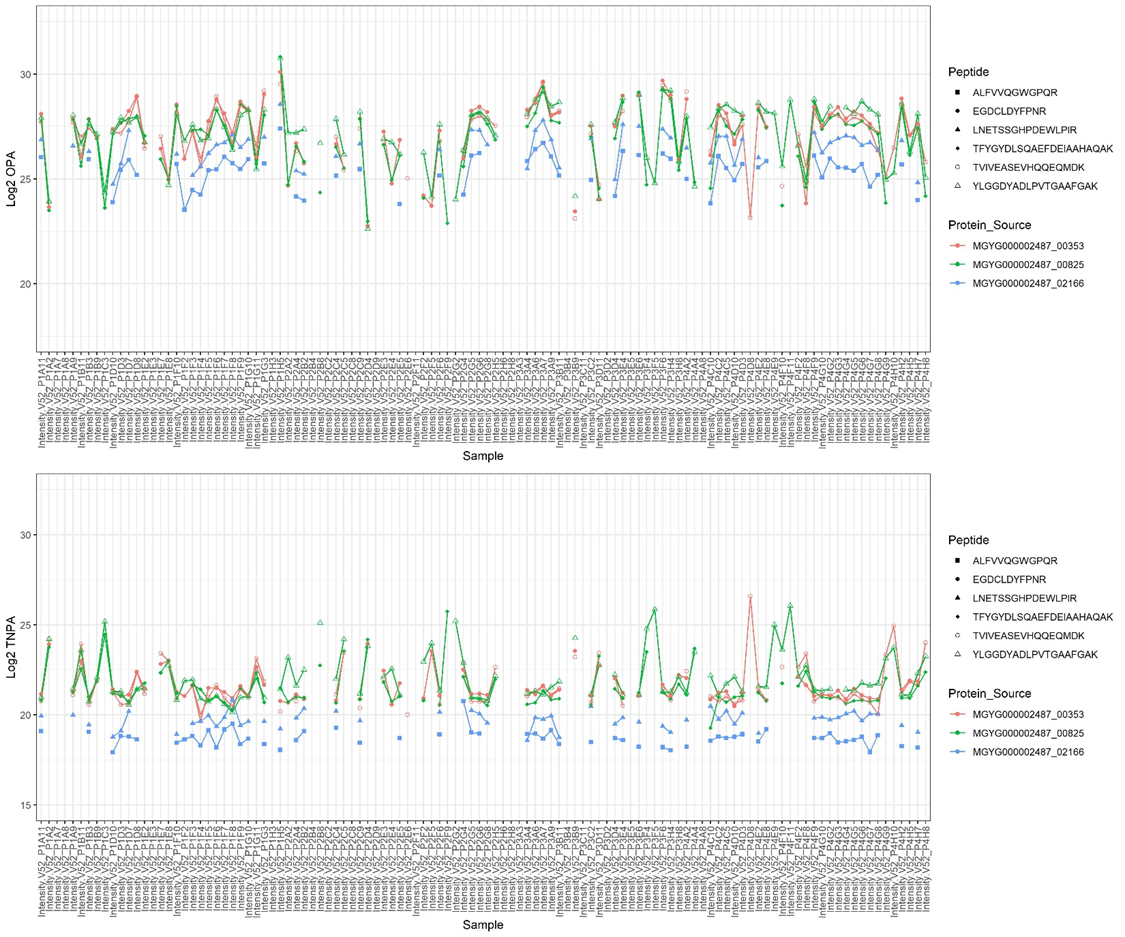

### Supplementary Figures 11.

**Changes in original peptide abundance (OPA, top panel) and taxa-normalized peptide abundance (TNPA, bottom panel) across different treatments for peptides from three proteins.** These proteins, originating from the same genome, have distinct functions: MGYG000002487_00353 (inosamine-phosphate amidinotransferase 1), MGYG000002487_00825 (putative protein), and MGYG000002487_02166 (putative dimethyl sulfoxide reductase chain YnfE).

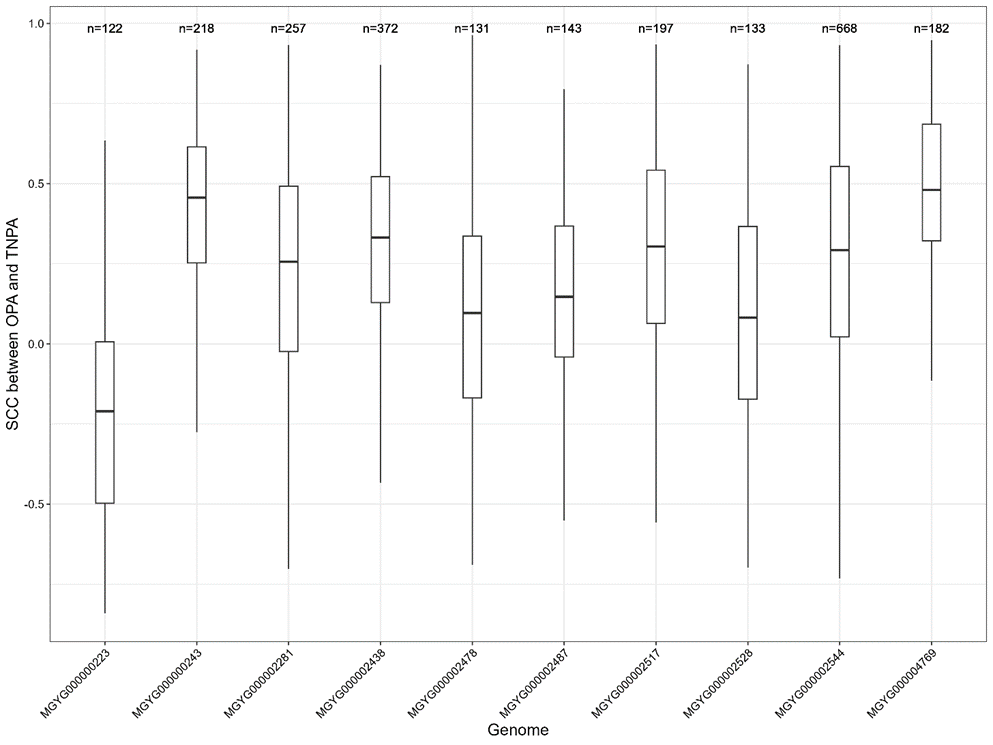

### Supplementary Figures 12.

**Distribution of SCC between OPA and TNPA for all peptides from each of the top 10 genomes.** n represents the number of peptides from each species.

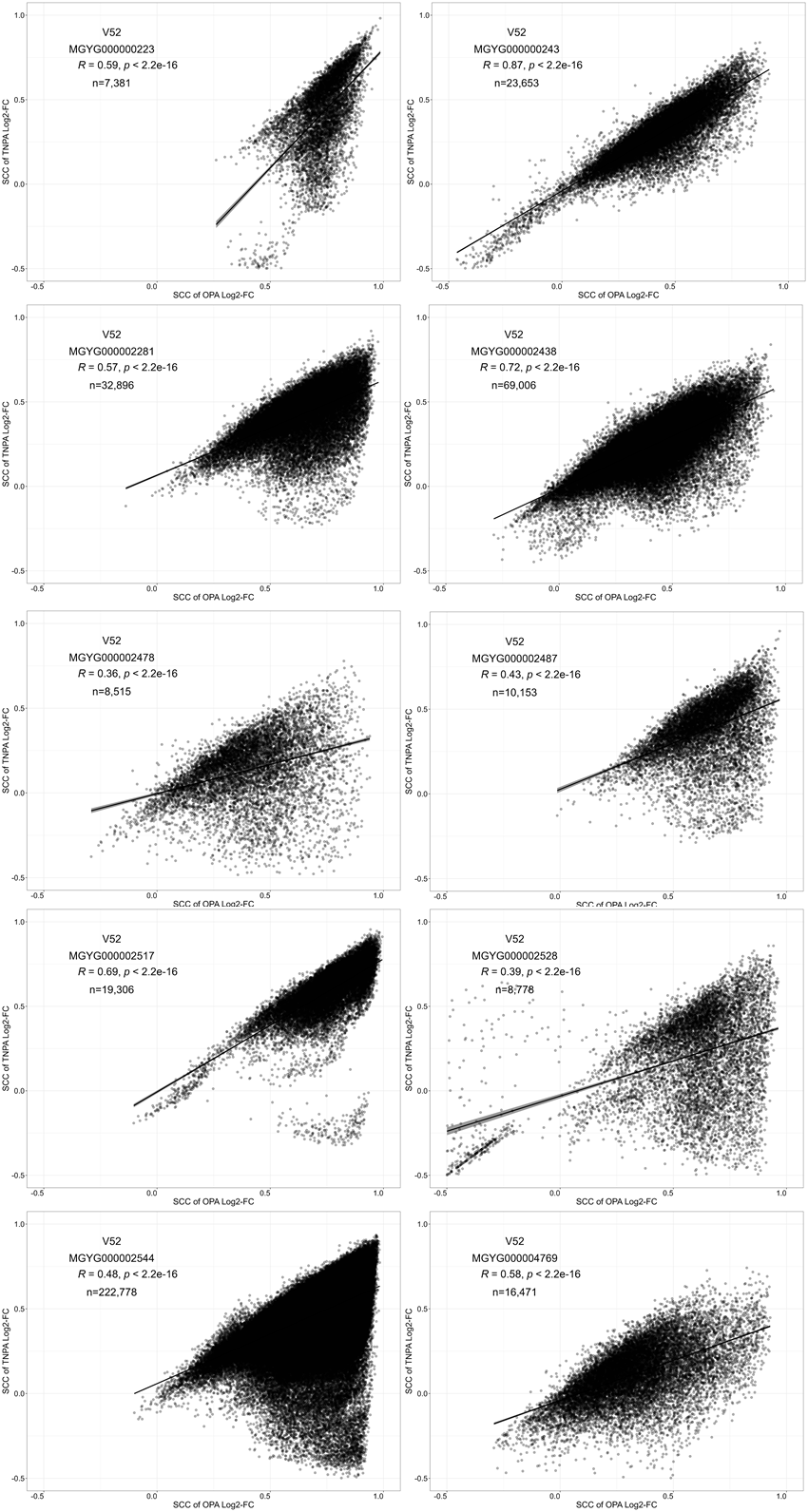

### Supplementary Figure 13.

**The correlation between the SCC of peptide abundance log2 FC based on TNPA and SCC of peptide abundance log2 FC based on OPA.**

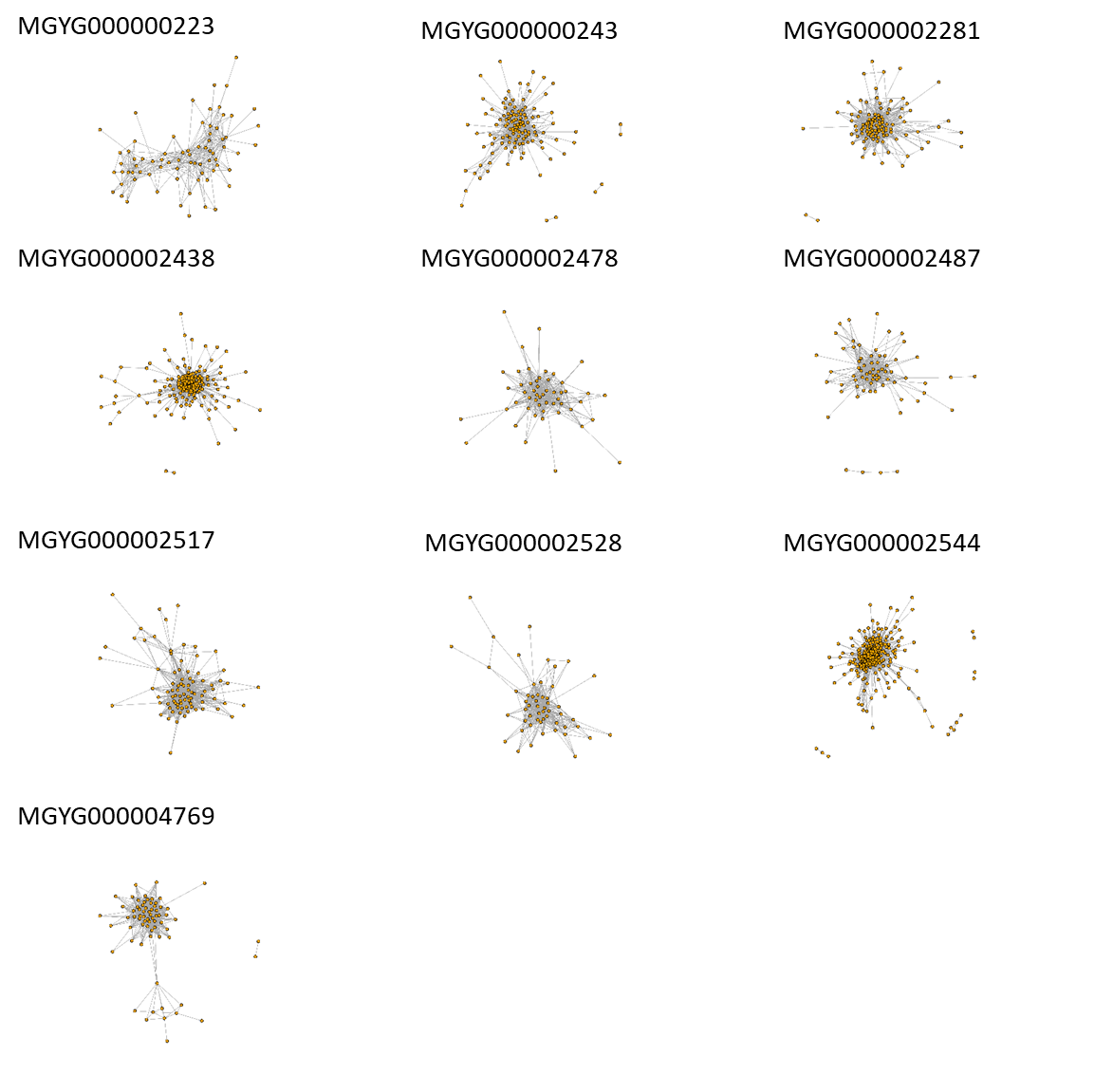

### Supplementary Figure 14.

**Single-species peptide abundance correlation networks based on OPA for the ten genomes with the most identified peptides from V52.** Each network was constructed using peptide pairs with the top 5% SCC from each species.

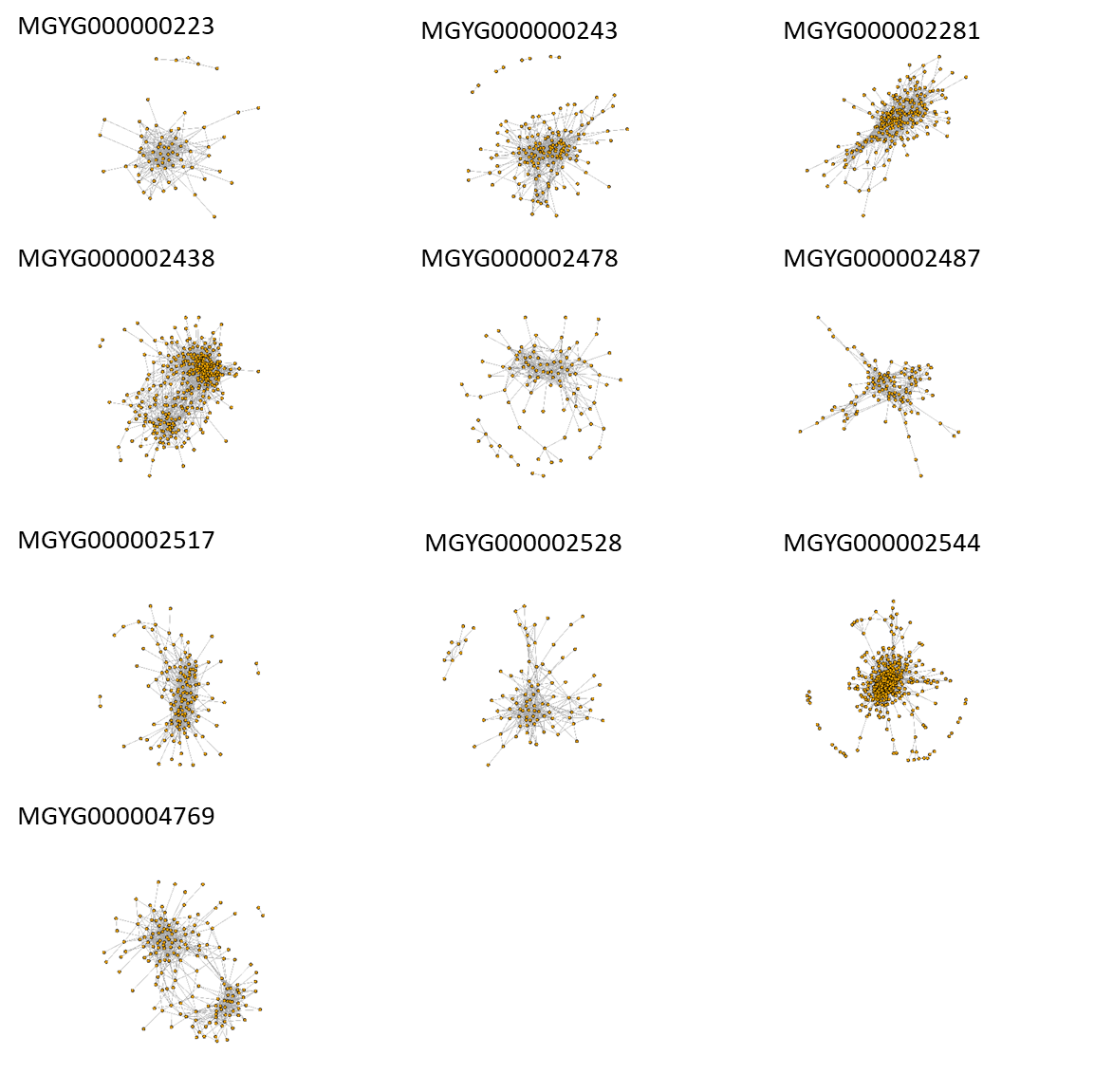

### Supplementary Figure 15.

**Single-species peptide abundance correlation networks based on TNPA for the ten genomes with the most identified peptides from V52.** Each network was constructed using peptide pairs with the top 5% SCC from each species.

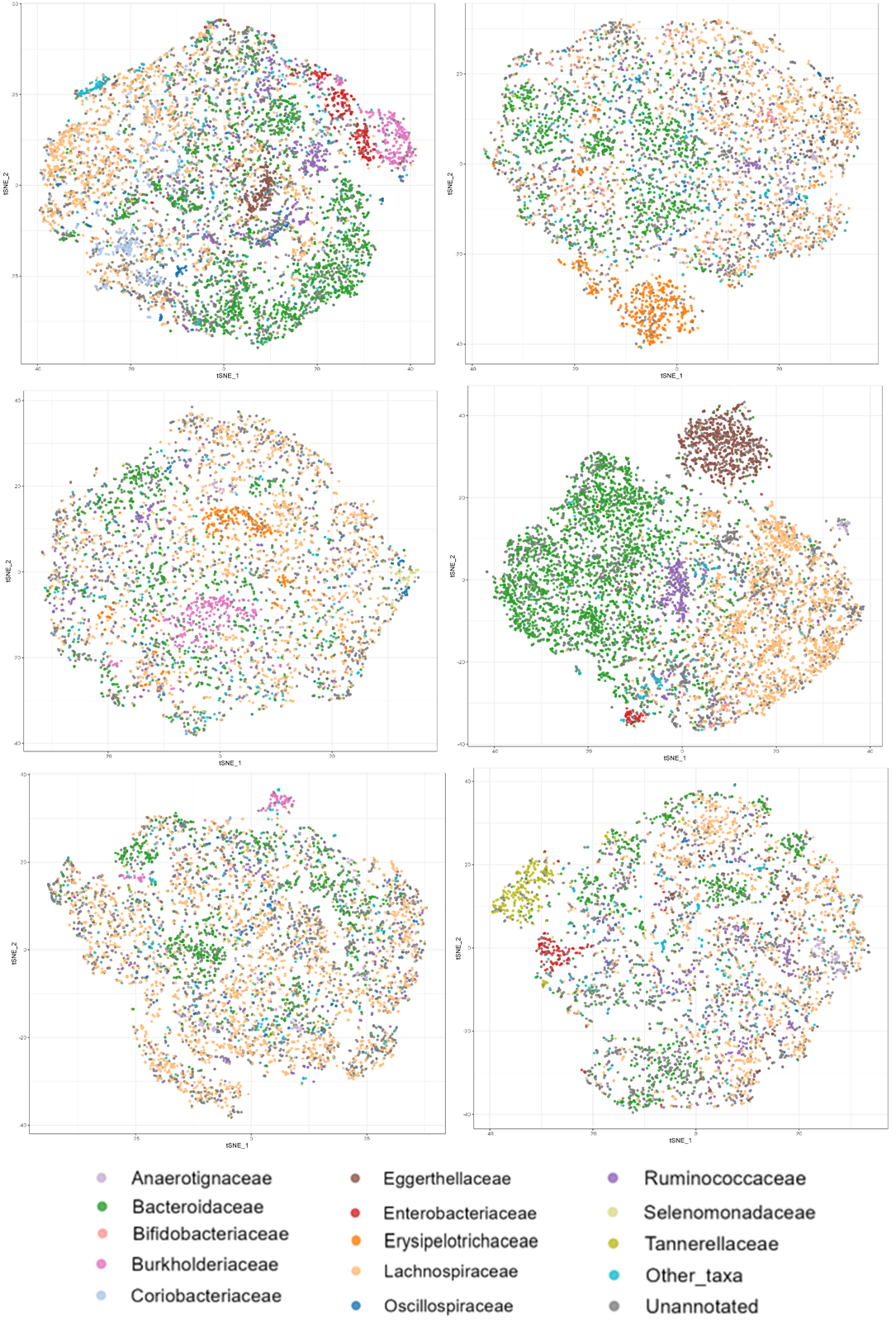

### Supplementary Figure 16.

**Global peptide correlation maps colored by peptide family-level taxonomic annotations from six individuals in another metaproteomics dataset (PXD012724).**
